# Geographical variation drives adaptive equilibrium of the *P. falciparum* sickle-associated mutations

**DOI:** 10.1101/2025.08.31.672853

**Authors:** Andre Python, Annie J. Forster, Peter Todd, Karamoko Niaré, Melissa D. Conrad, Lucas N. Amenga-Etego, Deus S. Ishengoma, Jianjun Lian, Baoli Liu, Yucheng Yan, Jingmin Liang, Muhammad M. Khan, Junkan Zheng, Jeffrey A. Bailey, David J. Conway, Alexander J. Mentzer, Alfred Amambua-Ngwa, Ellen M. Leffler, Thomas N. Williams, Gavin Band

## Abstract

The recent discovery of genetic mutations in *Plasmodium falciparum*—the most lethal malaria parasite—that enable it to overcome the protective effects of sickle cell trait, raises fundamental questions about the underlying biological and evolutionary interactions. Here we develop a geostatistical model to compare sickle haemoglobin genotype frequencies to the *Plasmodium falciparum* sickle-associated alleles across global populations, and find a robust association at multiple geographical scales, implying that sickle drives positive selection for these parasite mutations. A model of parasite evolution and an analysis of local haplotype patterns suggest that key features of these mutations – that they are polymorphic in all African populations and are mutually correlated despite lying in different genome regions - are caused by geographical variation in selection pressure, and that the alleles may have been maintained by balancing selection over timescales comparable to the age of the sickle mutation itself. The predicted impact of this host-parasite interaction on disease outcomes varies widely across populations, and functional data are needed to discover the biological mechanisms involved.

## Introduction

*Plasmodium falciparum* (*Pf*), the most lethal malaria parasite infecting humans, is a major selective force in human populations living in malaria-endemic regions. Exemplifying this, the sickle haemoglobin allele (HbS) is one of the best-known instances of balancing selection in the human genome, with the prevailing assumption being that the presence of *Pf* malaria in an area causes positive selection of HbS due to its protective effect against severe disease and death. This is balanced by the high mortality of HbS homozygotes due to sickle cell disease. Evidence for this hypothesis comes both from epidemiological estimates of the underlying effects^1^ and detailed geographical studies showing association of *Pf* prevalence with HbS allele frequencies globally^2,3^, and is the subject of continued investigation.

However, recent studies have shown that mutations in four distinct regions of the *Pf* genome strongly associate with both severe and mild infections of HbS-carrying individuals^4,5^. A natural hypothesis is that these mutations, which have been termed ‘*Plasmodium falciparum* sickle associated’ and here denoted *Pfsa+*, allow parasites to overcome the protective effects of HbS, and thus may be positively selected in HbS individuals. It is unclear, however, why the *Pfsa+* alleles have neither reached fixation nor been lost in African parasite populations and why they appear in strong linkage disequilibrium despite lying in different regions of the genome^4,5^. These features suggest complex opposing selection forces may be in operation. Understanding these forces may help explain how malaria parasites evolve to overcome naturally occurring resistance, and, potentially, make predictions about how this important interaction will evolve in future.

Here we contribute by analysing available geolocated HbS and *Pfsa+* genetic survey data covering malaria-endemic regions worldwide to quantify the association between HbS and *Pfsa+* frequencies at global and local scales. We use this to inform estimates of epidemiological parameters, including the underlying fitness effects of parasites given their genotypes and those of their hosts. Using a simulation of parasite evolution across the HbS frequency surface we then propose a model in which the *Pfsa* allele frequencies, and the between-locus linkage disequilibrium, are maintained by opposing and geographically varying selection pressures and as a result stay stable over time. An analysis of haplotypes at the loci also suggests long-standing maintenance of the alleles and, despite considerable uncertainties in the underlying parameters, suggests the mutations have been present for at least several millennia.

### Quantifying the global relationship between HbS and *Pfsa+* allele frequencies

We first quantified the relationship between HbS and *Pfsa+* allele frequencies in malaria-endemic populations worldwide. Our analysis is based on 1,098 geolocated HbS survey data points curated from multiple sources identified in a previous work^2^ and using a literature search (Table S1), and on *Pf* survey data comprising over 27,875 geolocated samples drawn from several published studies which used whole-genome sequencing^6,7^ or molecular inversion probe typing^8,9^ to assay genome wide genotypes (Fig. 1A and Table S2). We also generated new whole genome sequencing data for samples from Uganda (UCSF EppiCenter) and The Gambia (GAMCC study). A full description of our approach is given in Methods, but in brief, we first used the HbS survey data to estimate a global HbS allele frequency surface using a geostatistical modelling approach (posterior mean frequency shown in Fig. S1A and across Africa in Fig. 1B). This produced similar results to previously reported HbS frequency maps^2,10^ with minor differences due to modelling assumptions and the inclusion of additional data points (Fig. S2). To connect this to parasite data, we aggregated the estimated frequency of HbS genotypes, and those of parasite genotype counts from the *Pf* population-genetic survey data, in grid cells covering malaria-endemic regions worldwide (Fig. 1B; for the main analysis we used hexagons with a 1-degree long diagonal, or approximately 110 kilometres at the equator). For simplicity, in the analysis below we focus on the combined frequency of AS and SS genotypes in the population (denoted *f*_HbAS/SS_). We then developed a spatial regression model approach to estimate the relationship between *f*_HbAS/SS_ and *Pfsa+* genotype frequencies across grid cells (Fig. 1, C and D). Our approach uses a Bayesian framework throughout and quantifies statistical uncertainty by sampling from the full posterior distribution, allowing us to obtain estimates and statistical measures including a credible interval.

**Figure 1.**
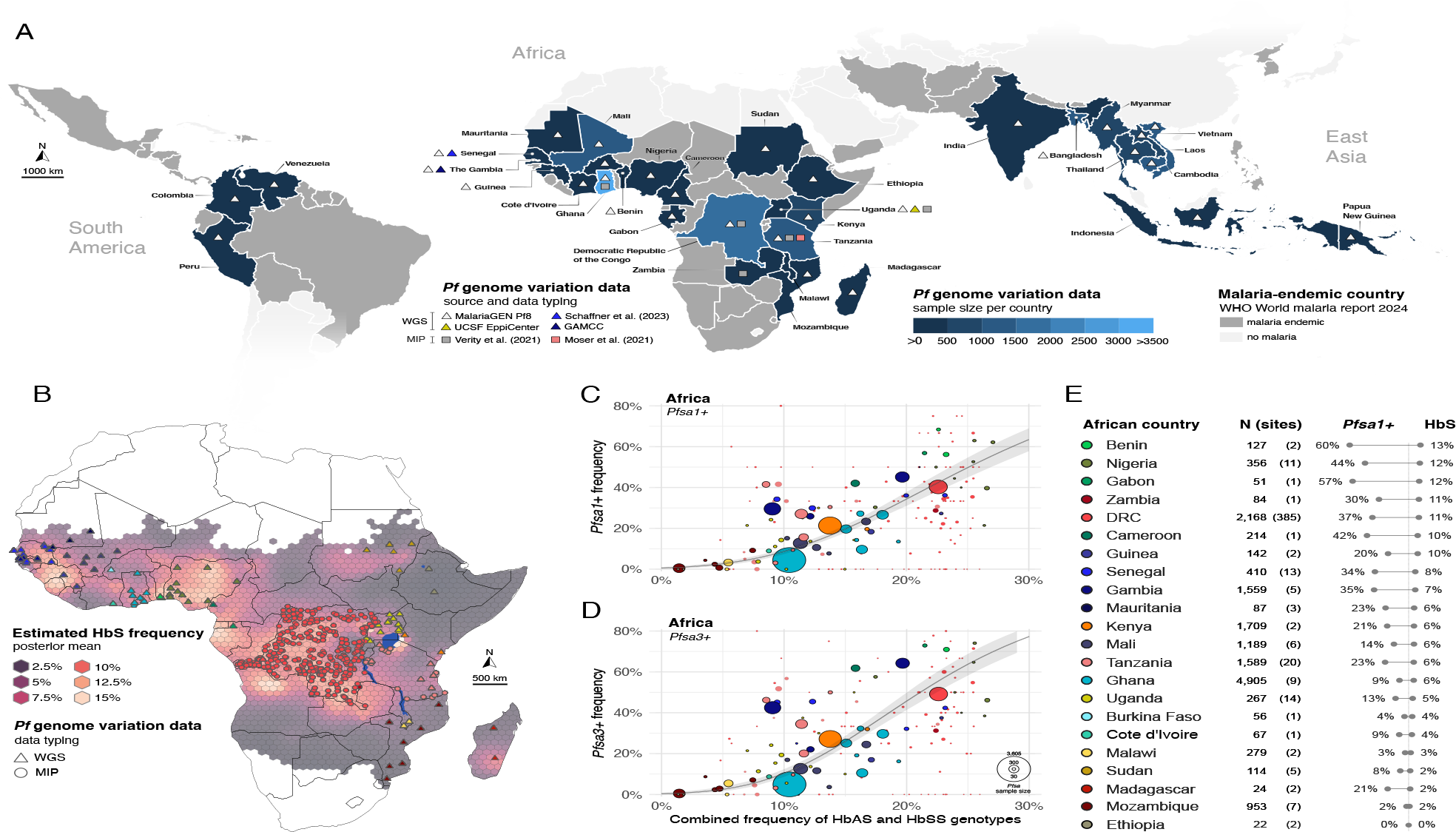
Statistical estimation of the HbS-*Pfsa* allele frequency relationship across African populations. **(A)** Countries containing *P. falciparum* (*Pf*) genome variation survey data (27,875 samples across 34 countries in Africa, South America, and East Asia) included in this manuscript. Countries without *Pf* data samples (dark grey areas) or assumed epidemic or malaria-free^11^ (light grey areas) are excluded from our analysis. (**B)** The estimated posterior mean frequency of the HbS allele across Africa (background colour) is illustrated, with the locations of *Pf* survey data from whole genome sequencing and molecular inversion probe typing illustrated as triangles and circles respectively. For illustration purposes, HbS allele frequencies are aggregated across hexagonal cells with a 1-degree long diagonal; only cells in malaria-endemic regions (determined using an estimate of the *Pf* parasite rate in children between 2 and 10 years old as observed in 2000^12^) are shown. A global HbS frequency map is illustrated in Fig. S1. (**C)** The relationship between HbS genotypes and *Pfsa1+* allele frequencies in African populations, estimated using a Bayesian spatial regression model (a modified Besag-York-Mollié^13^ model incorporating a generalised logit link function). Circles reflect the posterior mean estimate of the combined frequency of HbAS and HbSS genotypes (*f*_HbAS/SS_, x axis) and the estimated *Pfsa1+* frequency (y axis) in each hexagon. The circle size and colour reflect the *Pf* sample size and the country containing the *Pf* data points, as shown in the legend. Where grid cells overlap multiple countries, colouring reflects the country with the majority of data points. The black line reflects the mean regression model estimate of *Pfsa1+* frequency given *f*_HbAS/SS_ with uncertainty estimates (transparent grey lines) obtained from 10,000 samples of the joint posterior distribution. (**D)** Same as panel C but with *Pfsa3+* frequency on the y-axis. (**E)** Summary by African country of data collected at *Pf* survey points, with the sample size (N) and number of *Pf* sites along with average of *Pfsa1+* and HbS allele frequencies.

We applied this analysis to the three *Pf* genetic variants identified as the most strongly associated with HbS in the previous study of severe malaria cases^4,5^, namely *Pfsa1+* (defined as chr2:631,190 T>A with respect to Pf3D7_v3 reference assembly coordinates), *Pfsa2+* (chr2:814,288 C>T, or taken as chr2:814,329 A>G in MIP typing datasets^9^), and *Pfsa3+* (chr11:1,058,035 T>A). In addition, we analysed a fourth allele (*Pfsa4+*; chr4:1,121,472 T>A) which has been reported associated with HbS in a set of mild malaria cases in Ghana^4^. Details of the overall counts of these variants are given in Table S2.

Fig. 1C,D show the results of this analysis applied to *Pfsa1+* and *Pfsa3+* across African populations. The estimated relationship suggests that *Pfsa+* frequencies (*f_1+_* and *f_3+_*) are, on average, around 1.5 – 2 times as high as *f*_HbAS/SS_ in areas where the sickle allele is prevalent (e.g. *f_1+_* = 34% when *f*_HbAS/SS_ = 20%; 95%CI = 31% - 38%), but lower in areas where it is less common (e.g. *f_1+_* = 7% when *f*_HbAS/SS_ = 10% (6% - 9%)). This relationship also appears somewhat stronger for *Pfsa3+* than for *Pfsa1+*. To measure the overall strength of association we consider the difference in the estimated frequencies between *f*_HbAS/SS_ = 20% and *f*_HbAS/SS_ = 10%, which we denote by Δ*f_+_*. For *Pfsa1+* and *Pfsa3+*, Δ*f_1+_* = 26% (21-30%) and Δ*f_3+_* = 35% (29-41%) indicating strong positive relationships that are not plausibly caused by statistical sampling effects. We also confirmed that these results do not substantially depend on our underlying modelling choices (Figure S3 and Table S3).

The results above apply to African populations, but similar results were also obtained when fitting all global populations (Table S3). Moreover, we found that the positive association between *f*_HbAS/SS_ and *Pfsa+* frequencies was also apparent when restricting to western, or central and eastern African populations (Fig. 2) and indeed, a positive trend was observed within each individual country having a sufficient geographical spread of data to make estimates (Fig. S4) (although statistical resolution in many of these regions is limited).

**Figure 2.**
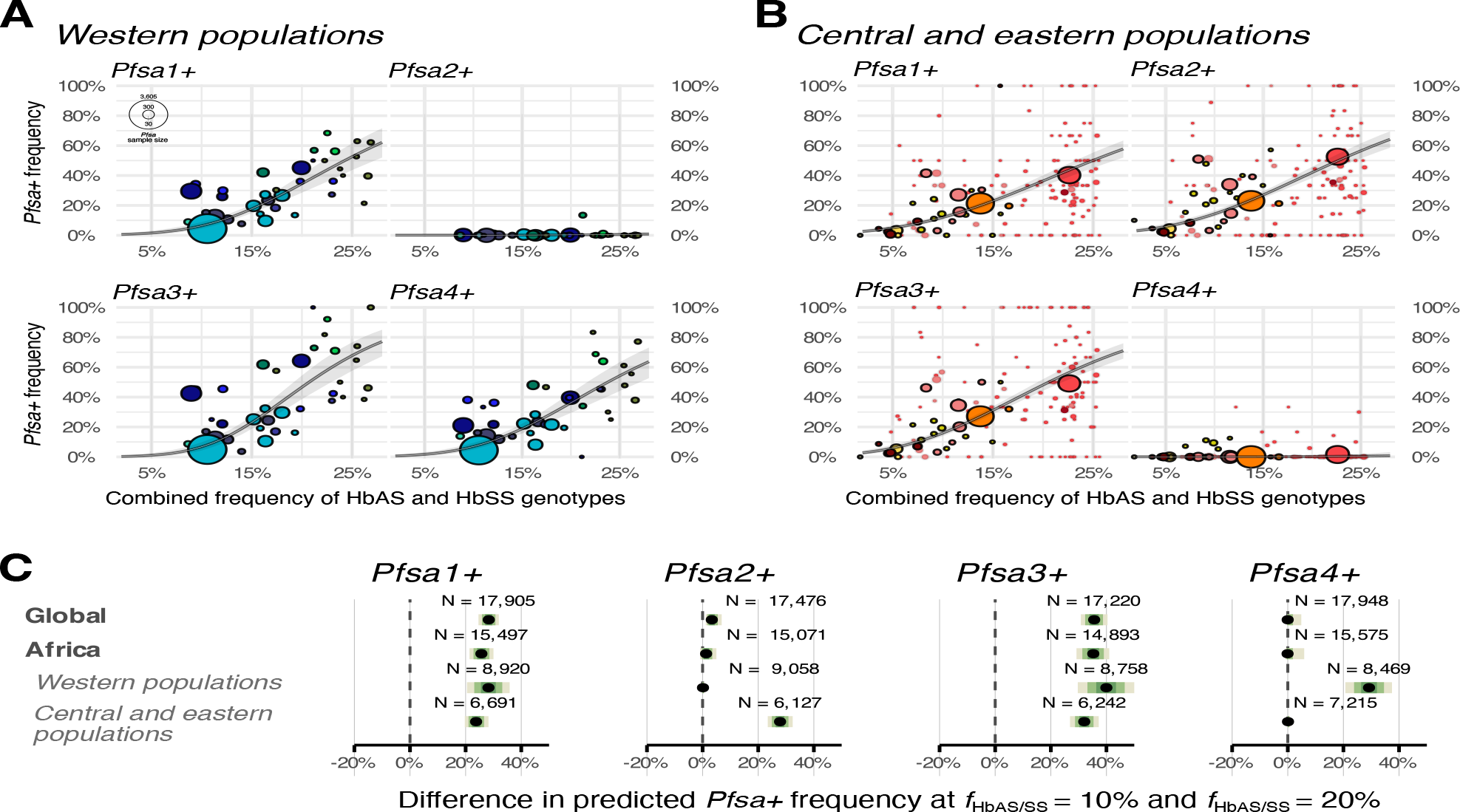
HbS-*Pfsa* relationship across spatial scales. **A-C.** Comparison of *Pfsa1-4+* and the combined frequency of HbAS and HbSS genotypes (*f*_HbAS/SS_) in (**A**) Western and (**B**) Central and eastern African populations. Each point represents a single grid cell with posterior mean estimate of *f*_HbAS/SS_ (x axis), *Pfsa+* frequency (y axis) and colouring as in Fig 1C,D. Black lines and grey ribbons depict the posterior mean regression model estimate of *Pfsa+* frequency and 95% credible intervals, similar to Fig. 1, C and D but estimated using the data only in those populations. For this analysis, the Western populations include Mauritania, Senegal, Gambia, Guinea, Mali, Burkina Faso, Cote d’Ivoire, Ghana, Benin, Nigeria, Cameroon, and Gabon, and Central and eastern populations include Democratic Republic of Congo, Ethiopia, Sudan, Kenya, Uganda, Malawi, Zambia, Mozambique, Tanzania, and Madagascar. These divisions are chosen to correspond to the geographic areas of Africa where *Pfsa2+* and *Pfsa4+* are broadly prevalent; exceptions to this are visible in panels A,B and noted in main text. (**C)** Summary of the estimated regression slope (taken as the difference in predicted *Pfsa+* frequency between *f*_HbAS/SS_ = 10% and *f*_HbAS/SS_ = 20%; denoted Δ*f_+_* in main text) across 10,000 draws from the posterior distribution, in global regions, Africa, and in each African region. Black points represent the posterior median estimate with shades of green indicating the 95%, 80%, and 50% posterior intervals.

Analysis of the *Pfsa2+* and *Pfsa4+* alleles is more complicated because, as previously observed^4,5^, the *Pfsa2+* allele is only prevalent in eastern African regions and the *Pfsa4+* mutation is primarily present in west and central African regions. However, in both cases the allele frequencies show similar strong association with HbS where the allele is present (Δ*f_2+_* = 27% (23-32%) across DRC and east African populations; Δ*f_4+_* = 29% (21-38%) in west and central African populations). In particular this analysis provides additional support for a relationship between *Pfsa4+* and HbS beyond the limited association data reported so far^4^. It is however worth noting that these alleles are also present at some sites outside these overall patterns (e.g. *Pfsa2+* observed in >10% of samples in Gabon (N=52) in west Africa; *Pfsa4+* observed in >1% of samples in Uganda (N=275), DRC (N=2,403), and Zambia (N=92) in east Africa). Moreover, *Pfsa2+, Pfsa3+*, and *Pfsa 4+* are also present in Asian and especially South American populations (Table S2); the latter may reflect the likely origin of South American *P. falciparum* parasites in the transatlantic slave trade^14^ and the fact that the majority of complete South American *P. falciparum* genomes in the MalariaGEN dataset were sampled from populations with predominantly African ancestry^15^.

The above results demonstrate that the frequencies of all four *Pfsa+* alleles are associated with that of HbAS and HbSS genotypes across Africa or within African regions. Moreover, the alleles are also in strong linkage disequilibrium (LD) (i.e. the *Pfsa+* alleles tend to be observed together in the same parasites) across all geographic regions (Fig. S5). Together with published data showing elevated rates of *Pfsa+* parasites among severe and mild infections of HbAS/SS individuals^4,5^ the results strongly support the hypothesis that *Pfsa+* alleles are positively selected for by HbS. However, this raises the question of why the mutations have not reached fixation in any location, as well as by what mechanism the linkage disequilibrium is maintained.

### Maintenance of the *Pfsa* polymorphisms by geographically varying selection

To explore possible evolutionary scenarios we investigated population-genetic models of parasite evolution taking place over the HbS genotype frequency map across Africa, that capture three main factors that could play a role. First, we modelled parasite fitness effects (interpreted as the probability that a bite by an infected mosquito will lead to a successful infection that is transmissible to future generations) that depend on both host and parasite genotypes. Second, we modelled the effects of geographical variation in HbS genotype frequencies – and therefore variation in selection pressure on *Pfsa* genotypes – by imposing a limited geographical range on mosquito biting distance in each generation. And third, we modelled the effects of transmission on breakdown of linkage disequilibrium (restricting to two modelled *Pfsa* loci for simplicity) by allowing for meiosis during a proportion of transmissions.

Although the underlying fitness parameters are not known, plausible constraints are provided by the above analysis of geographical variation (Fig. 3A and Fig. S6) and by current estimates of relative risk of infection conferred by HbS across *Pfsa* genotypes in severe infections^5^. Accordingly we focus here on models in which the protection afforded by HbS is strong against *Pfsa-* genotype parasites, but weak or negligible against *Pfsa+* genotype parasites. Moreover, since *Pfsa+* genotypes are also observed in substantial numbers of infections of HbAA individuals we also assume that the overall fitness cost of *Pfsa+* genotypes must be relatively weak. A specific set of fitness parameters meeting these criteria is shown in Fig. 3B.

**Figure 3.**
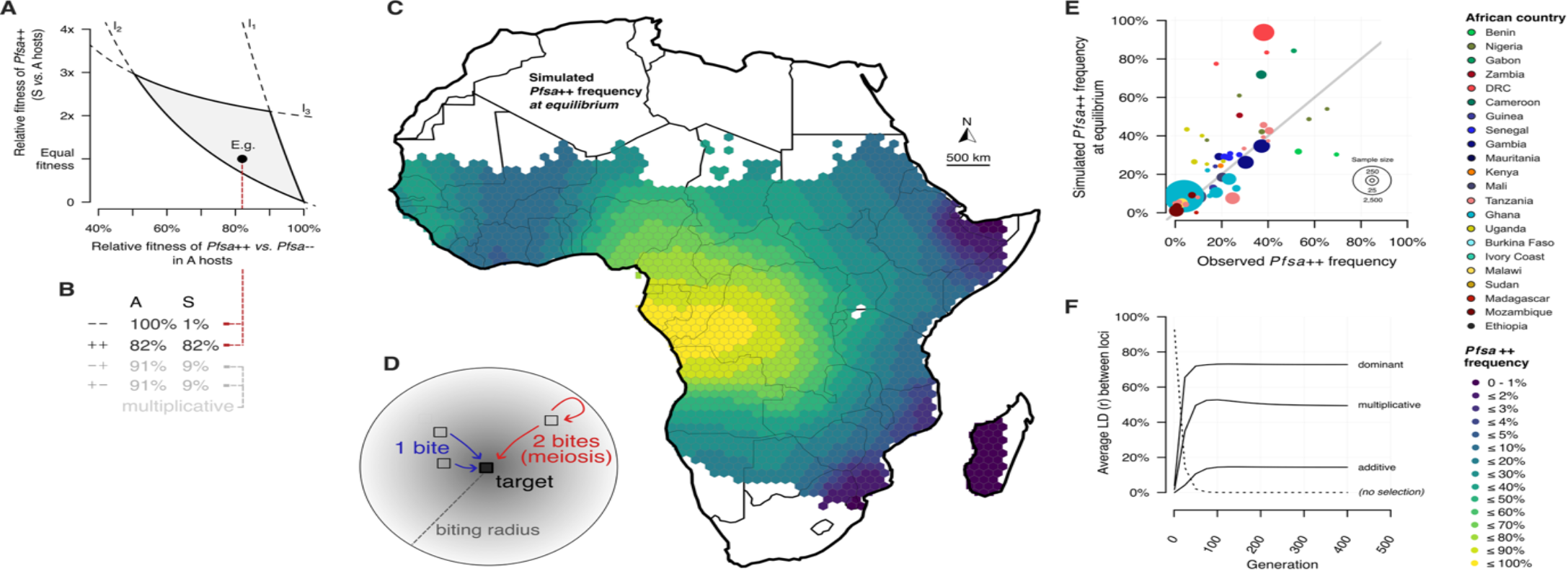
Stable *Pfsa+* equilibrium patterns arise from geographical variation in HbS frequency. **(A)** Constraints on parasite fitnesses implied by the geographical analysis in Fig. 1C,D, under the assumptions that ++ parasites are not positively selected at *f*_HbAS/SS_< 5% and not negatively selected at *f*_HbAS/SS_> 25% (lines *l*_1_ and *l*_2_), and that HbS does not confer increased overall parasite fitness (line *l*_3_) (Methods). In the figure we assume that -- parasites have a 100-fold fitness cost HbS genotype individuals (γ‐ ‐ *s*= 1%); other choices are shown in Fig. S6. (**B**) Example fitness parameters across four parasite genotypes matching the constraints in panel A, symmetric between the alleles and approximately multiplicative within A or S hosts, such that -+ and +- genotypes have intermediate fitnesses. (**C**) Visualisation of parasite allele frequencies obtained from a discrete-time model of evolution over malaria-endemic regions of Africa. Colours range from dark blue (low) to yellow (high) allele frequencies as in the legend. Evolution proceeds in each timepoint by replacing the parasite population at each location with the content of infected mosquitoes, which sample their parasites by from the local area (depicted in panel D). The rate of success of each infection (i.e. whether it is allowed to contribute to the updated local population frequencies) depends on both the host and parasite genotypes, according to the relative fitnesses shown in panel B. In the simulation shown, sampling locations are weighted by an estimate of local *Pf* prevalence^12^ and inversely with distance to the target location. To model breakdown of linkage disequilibrium, a proportion *m* = 15% of mosquitoes at each location are assumed to sample two parasites, which undergo meiosis. After approximately 200 generations, the model reaches a stable equilibrium at which the *Pfsa* allele frequencies no longer change and the loci are polymorphic at all geographic locations. **(E)** Comparison of the simulated frequency of the ++ allele (*y axis*) to the observed frequency of *Pfsa1+* / *Pfsa3+* parasites (*x axis*). Points are sized for sample size and coloured by country as in Fig. 1E. **(F)** Evolution of linkage disequilibrium (LD; measured as the correlation (r) between the + alleles at the two loci (*y axis*), averaged over sites across generations (*x axis*). Solid lines indicate three different sets of fitness costs for the +- and -+ genotypes. The multiplicative model is shown in the table while additive and dominant models have, respectively, higher and lower relative fitnesses for +- and -+ genotypes compared to the multiplicative model. The dashed line indicates the behaviour of the model starting from 100% correlation if there is no selection (all relative fitnesses = 1).

Figure 3C shows the results of parasite evolution under this setting (implemented using the WebGPU framework; see Data and materials availability). We found that these models display a general convergence behaviour, limiting to an equilibrium pattern of allele frequencies that is maintained over time. The equilibrium obtained depends on the fitness parameters, but for parameters consistent with those motivated above it maintains polymorphism at both *Pfsa* loci across all geographic areas. Consistent with Fig 3A, this behaviour appears when the fitness cost of parasites with both *Pfsa+* alleles (++) is modest (for example, a relative fitness (γ_++_) of approximately 80%) and the fitness cost against -- parasites in S individuals is substantial (e.g. γ‐ ‐ ≈ 1 − 10%) (Fig. 3B). For some parameter choices, it produces frequency patterns at equilibrium that are reasonably consistent with observed data (Fig. 3E).

Moreover, a further feature of these models is that they generate strong between-locus linkage disequilibrium patterns across loci (Fig. 3F). This occurs even when no LD is present at the start of the simulation and arises due to the joint selection acting at the two loci - even when the rate (*m*) at which mosquitoes sample multiple parasites which undergo meiosis is sufficient to otherwise break down LD relatively rapidly in the absence of selection (e.g. *m* = 15% in Fig. 3). The level of LD obtained depends both on this rate and on the relevant fitness of the - + and +- genotype combinations, but we found it can be substantial even when there is no epistasis in either A or S individuals (e.g. under additive or multiplicative fitness models; Fig. 3F with details in materials and methods). We emphasise, however, that our analysis does not rule out a level of between-locus epistasis, which may be considered plausible given the high levels of LD observed.

This analysis therefore suggests a plausible mechanism that could generate patterns of intermediate allele frequencies, and of long-range LD, that remain stable over time and are qualitatively similar to those observed in current populations. We emphasise that this behaviour arises from the geographical variation in selection in the model; similar behaviour holds for single-locus models but not for selection models in homogeneous populations (although other types of negative frequency-dependent selection are possible). These results also closely align with earlier theoretical work demonstrating conditions under which similar behaviour holds in models of discrete demes or continuously varying populations with migration^16-20^.

### Local haplotype variation at *Pfsa1* and *Pfsa3* indicate long-term balancing selection

The above analysis suggests a mechanism by which polymorphism of the *Pfsa* variants may have been maintained over long time periods. If so, this may have left signatures in the patterns of genetic variation along chromosomes close to the loci. To assess this, we used a version of the MalariaGEN Pf7 dataset^21^ filtered for population-genetic analyses by focussing on unmixed infections from African populations and to confidently-called genetic variants. We further polarised variants by using data from gorilla- and chimpanzee-infecting malaria species (*P. praefalciparum* and *P. reichenowi*), which are closely phylogenetically related to *P. falciparum^22^*, to identify the ancestral and non-ancestral (i.e. derived) allele at each of (N=105,000) sites across the core genome.

Fig. 4 illustrates the chromosomes (i.e. haplotypes) of parasites in a 10kb region centred on *Pfsa1* in this dataset, together with a genealogical tree which we estimated using RELATE^23^. In the following we refer to the haplotypes as *Pfsa1+* or *Pfsa1-* according to whether they carry the *Pfsa+* mutation at the lead SNP (which is indicated with an inverted triangle in Fig. 4). A total of 12 additional common (frequency > 10%) derived mutations were observed that are carried almost exclusively on *Pfsa+* haplotypes, across a region of 5kb centred on the lead mutation (Fig. 4C). This pattern of allele sharing therefore indicates that the *Pfsa1+* haplotypes originated in a common ancestral parasite, DNA from which has been inherited by all African populations (Fig 4A and B).

**Figure 4.**
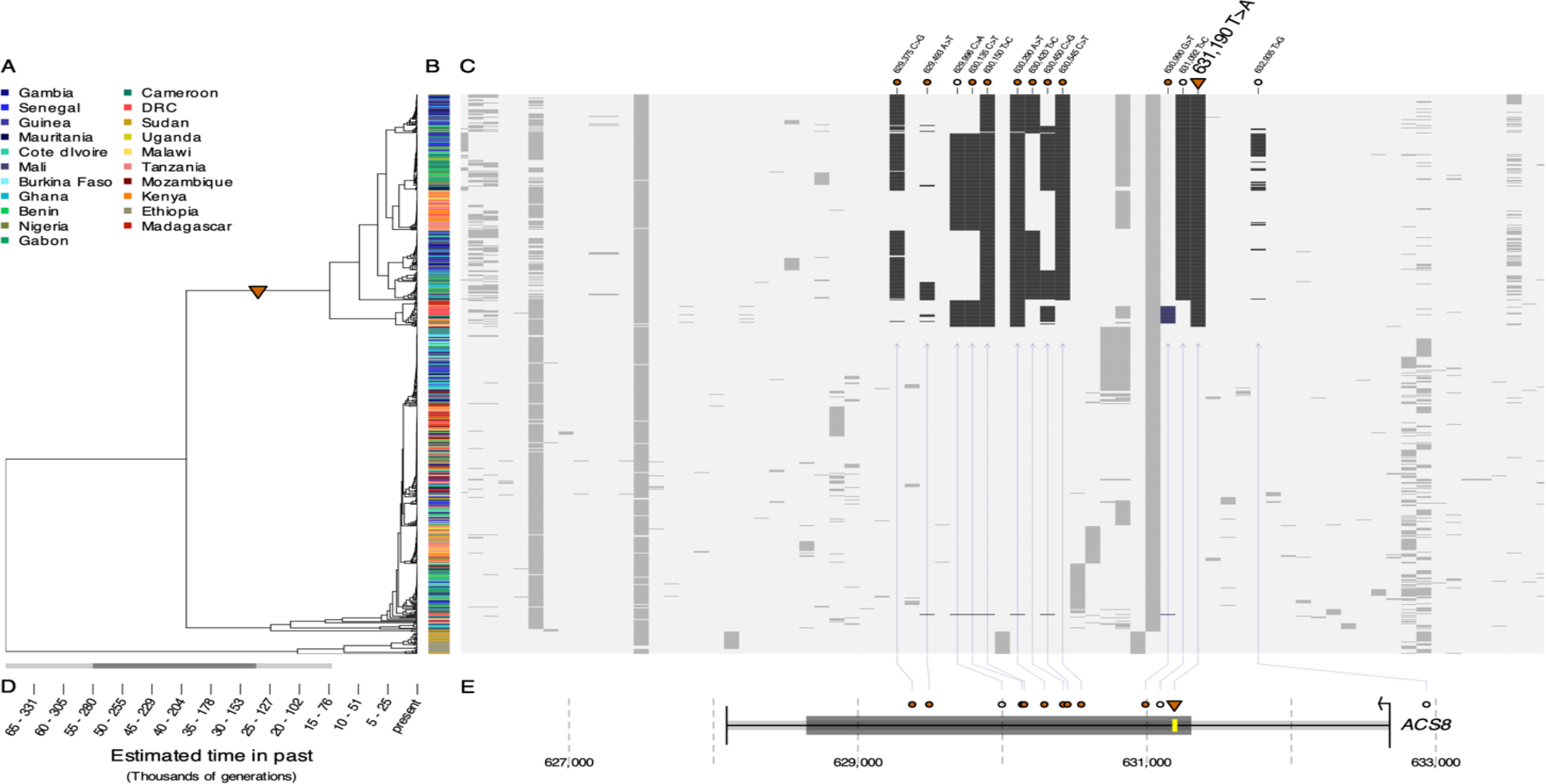
Haplotype patterns suggest long-term maintenance of *Pfsa* polymorphism at the *Pfsa1* locus. (**A**) Genealogical tree estimated at the *Pfsa1* lead SNP (chr2:631,190 T>A) using RELATE^23^, based on N=4,788 non-mixed parasite samples from African populations in the MalariaGEN Pf7 resource. For visualisation purposes, the tree is sub-sampled to a maximum of 20 samples per country and per *Pfsa1*+ genotype. The inverted triangle indicates the branch corresponding to the lead *Pfsa1+* allele. (**B)** Country of ascertainment of each infection in the tree (colours, as in legend). (**C**) Pattern of ancestral (*light grey*) and derived (i.e. non-ancestral) alleles (*dark grey*) carried by each infection (rows), using the same samples and ordering as shown in the tree in panel A. Details of the location of common derived mutations linked to the *Pfsa*+ haplotypes (defined here as appearing in at least 10 *Pfsa+* haplotypes and fewer than 10 *Pfsa-* haplotypes) are shown, with non-synonymous mutations denoted in orange circles and synonymous or intragenic as open circles. Parasites from Sudan and Ethiopia carry a distinct haplotype which carries a different set of derived mutations (bottom rows of panel B) and is assigned to an ancestral branch. (**D**) Branch length estimates for the tree in A assuming a mutation rate of either 6.3162 × 10^./^ or 3.21552 × 10^.0^ per transmission cycle^22^. Above, bounds on the estimated age of the *Pfsa1+* allele are shown (95% interval, light grey, and 50% interval, dark grey). (**E**) Detail of the location of the highlighted mutations within the ACS8 gene. The yellow rectangle indicates the location of a predicted PEXEL motif^24^ which is by the *Pfsa1*+ lead mutation (KAIYE → KAIFE).

Patterns of derived mutations of the form seen in Fig. 4 are a possible signature of balancing selection, under which long-term maintenance of polymorphism at a locus can lead to buildup of nearby co-segregating mutations^25,26^. To test for this at *Pfsa1*, we computed metrics reflecting different modes of selection including recent positive selection (iHS, iHH12) and balancing selection (Tajima’s D, Beta, Dango^27^), as well as the proportion of nucleotide variation between the *Pfsa+* and *Pfsa-* alleles. We compared these metrics to frequency-matched variants genome-wide, working separately in each population, and used this to compute empirical P-values (Table S4). We found little evidence for a recent selective sweep of the *Pfsa1+* haplotype in any African population (e.g. *iHS* < 1.8 standard deviations from zero; empirical P-value 0.06-0.86; computed across variants within 1% frequency of that of the lead SNP in each population; after excluding variants near known drug resistance, hypervariable and antigenic genes as described in materials and methods). This includes populations where the allele frequency is especially high (e.g. Cameroon, *iHS* = −0.61; DRC, *iHS* = 0.65). The iHH12 metric, which is more suitable for the detection of positive selection on standing variation, similarly did not reach unusual values (normalised *iHH*12 ≤ 1.64) although we noted the statistic was positive in all populations. By contrast, metrics reflecting balancing selection showed empirically extreme values in several populations (e.g. Beta = 2.8 – 63.6 with empirical P-values = 0.006 - 0.13; Dango = 4.0 – 6.4 with empirical P-values = 0.01 – 0.09; computed across a 5kb window centred at each variant). A particularly clear signal was seen for a metric reflecting allele age (the average number of mutations separating *Pfsa+* and *Pfsa-* haplotypes, as a proportion of total nucleotide diversity (π_b*etween*_/π*_total_* = 37 – 85% with empirical P-values = 0.006 – 0.07 and < 0.05 in 9 of 10 populations).

These analyses therefore provide evidence that *Pfsa1+* is older than most other variants at similar frequency across the genome and potentially under balancing selection. An allele age estimate using previously computed upper and lower estimates for the core genome mutation rate applicable between transmissions^22^ (i.e. 6 to 32 × 10^09^ mutations per site per transmission cycle) suggests that *Pfsa1+* may have originated between 15,000 and 300,000 transmissions ago (Fig. 4D and Fig S6). Although there are further uncertainties involved in translating this to a duration in years, this likely indicates an origin at least several millennia ago.

We also observed similar results at the *Pfsa3* locus (Figure S), although we noted that the most compatible results were obtained when focussing on the chr11:1,047,437 T>C mutation (which had the second highest level of evidence for association with HbS in the previous analysis of severe cases^5^). We caution that this locus also contains structural variation, which may be linked to the *Pfsa+* allele^5^. The region-specific *Pfsa2+* and *Pfsa4+* alleles both appear to have originated more recently.

We note that while the origin of the HbS allele is still a matter of research, current analyses suggest it originated between 13,000 and 70,000 years ago in ancestors of modern-day agriculturalist populations^28,29^, and may have reached a balanced frequency within a few thousand years. Therefore, these results are compatible with a possible origin of the *Pfsa* alleles arising as an evolutionary response to the spread of protection due to HbS.

## Discussion

The HbS allele, which encodes sickle haemoglobin, is one of the best-known examples of balancing selection in the human genome. Its frequency is maintained by the protection it provides against malaria in endemic populations, and by the fitness cost associated with sickle cell anaemia in homozygotes. The combined evidence of the geospatial modelling, simulation and genetic analyses presented here suggests that the *P. falciparum* sickle-associated alleles, which appear to confer on parasites the ability to overcome the protection due to HbS, are also under a form of balancing selection. Our analysis also suggests a mechanism by which this may have arisen, namely that this is a natural consequence of the geographic variation in selection pressure, that is, variation in the prevalence of HbS itself.

A recent analysis of ancient and modern infections from the Gambia^30^ has identified a substantial increase in *Pfsa1+* and *Pfsa3+* mutations over the last 60 years from around 10% frequency in the late 1960s to the present-day frequency of around 50-70%. There is also some evidence of recent frequency changes in other populations (Fig. S9 and S10), although our results show that this has not been accompanied by strong haplotypic signals of recent positive selection. Given our results, it seems most plausible that these observations represent frequency changes of haplotypes that pre-existed in these populations, that is, selection – or genetic drift – operating on standing variation. In the context of our model, this could occur due to fitness alterations caused by changes to the environment in which parasites evolve, or due to changes in parasite populations which have occurred recently due to interventions. In particular, it is notable that the *Pfsa* loci are in positive linkage disequilibrium with mutations that confer chloroquine resistance in many areas (Table S5), which have seen even more striking changes over this time period.

The presence of selection may be surprising because HbS has generally been thought not to have a strong effect against asymptomatic infections, which contribute a substantial portion of transmissions. However, our results suggest the effect of HbS on parasite fitness – just as on severe outcomes – must indeed depend strongly on the parasite genotype. *Pfsa+* parasites also appear to suffer at least some fitness cost relative to *Pfsa-* parasites in HbAA individuals. It is likely that the parasites are less well adjusted to growth in HbAA erythrocytes, but previous work has also linked HbS (and the neighbouring mutation HbC) to altered rates of gametocytogenesis and transmission^31,32^, and it is therefore also possible that fitness costs may arise from a form of reduced transmissibility. Studies that compare parasites within blood stages and across the transmission cycle will be needed to resolve the underlying biological effects.

Ultimately the data outlined here and previously^4,5^ may be important because they suggest that for individuals with HbS genotypes, the risk of malaria infection and disease will vary considerably across geographical regions in Africa. In regions where *Pfsa+* alleles are at high frequency, HbS may provide little if any protection. We caution however that the full extent of parasite genetic loci which interact with the protective effect of HbS may not yet be known. Given the signals of longstanding maintenance of the alleles, and the geographically localised nature already observed for the *Pfsa2* and *Pfsa4* loci, a natural question is whether parasite populations have or are evolving other mutations that reduce the associated fitness costs. Current studies of host-parasite genetic association^4,5^ are underpowered to detect such alleles, but future parasite genome-wide association studies which increase the overall sample size and/or meta-analyse across populations may provide new information. Since the pattern of HbS frequencies is also likely to be changing over time, albeit at a lower rate, the current observations may be best thought of as a snapshot of a longer dynamic process of co-evolution between these host and parasite alleles.

## Acknowledgements

The authors would like to thank Jonathan Parr, Jon Juliano, Robert Verity and Bryan Greenhouse for assistance identifying relevant data and comments on the work. This publication uses data from MalariaGEN Pf7^21^ and MalariaGEN Pf8^6^, from samples collected under Demographic Health Surveillance Systems in the Democratic Republic of the Congo^8^ and Tanzania^9^, and from samples collected in Senegal in 2019^7^. This paper also uses previously unpublished parasite genetic data from Uganda and The Gambia for which sample collections were previously described^33-35^.

## Author contributions

Conceptualization: A.P., D.J.C., E.M.L., T.N.W., G.B. Methodology and formal analysis: A.P., J.L., A.J.F., G.B. Investigation: A.P., A.J.F., J.L., B.L., Y.Y., J.L., M.M.K, G.B. Visualization and software: A.P., P.T., G.B. Funding acquisition: A.P., B.L., D.J.C., T.N.W., G.B. Project administration: B.L. Supervision: A.P., A.J.M., G.B. Resources: A.J.F., K.N., M.D.C., L.N.E-E., D.S.I., J.A.B., A.A-N. Writing – original draft: A.P., G.B. Writing – review & editing: All authors.

## Data and materials availability

Analysis scripts and a Snakemake^36^ pipeline to reproduce the results of our analysis will be made available via GitHub. Data from previously published component datasets and relevant URLs is detailed in tables S1 and S2. GAMCC study data will be deposited in a public repository (accession tbc). Uganda study data will be deposited in a public repository (accession tbc.). The simulation of parasite evolution over the HbS frequency map can be found at https://www.chg.ox.ac.uk/~gav/projects/pfsa/simulation/.

## Supplementary Materials

Methods, supplementary text and supplementary figures can be found in the enclosed document “Methods and supplementary information.docx”.

Supplementary data is available in the file “Supplementary Tables.xlsx”.

