## Supplementary material for "Geographical variation drives adaptive equilibrium of the *P. falciparum* sickle-associated mutations": Methods and Supplementary Information

### Materials and Methods

#### 1.1 Ethical approvals

Ethical approval for *Pf* sample collections and sequencing in Uganda was obtained from the Makerere University Research and Ethics Committee, the Uganda National Council for Science and Technology, and the University of California, San Francisco, Human Research Protection Program. Ethical approval for re-analysis of DNA from samples collected in The Gambia (GAMCC study) was given by the The Gambia Government / MRC Joint Ethics Committee (26354) and OxTREC (509-22)

#### 1.2 Collation of sickle haemoglobin survey data

We gathered data on sickle haemoglobin prevalence from two sources. First, we used results from studies published between 1950 and October 2009, gathered by a previously conducted literature search[1] and downloaded on 8th October 2022 using the R package MALARIAATLAS[2]. This data included 1,286 geolocated records from 473 source publications representing 1,156 distinct locations globally. As in a published re-analysis of this data[3] we retained all studies where counts for HbAA and HbAS genotypes were provided (totalling 1,041 records); four of records were additionally removed based on inspection of the data. We then conducted a further literature search to identify studies published in the period November 2009 to November 2024. Specifically, we used PubMed, Scopus, and Web of Science and search terms "sickle cell", "haemoglobin S", "hemoglobin S" and "Hb S" with index dates from 20th October 2009 to 3rd November 2024, and where the publication contained genotyping or blood typing informative of local HbS prevalence; a total of 205 publications were obtained. We excluded studies that did not have sufficiently accurate georeferencing, requiring spatial accuracy to at least ADM-2, which corresponds to second-order administrative subdivisions within each country. As ADM-2 varies considerably in size between countries, we also required that the size of the reported area was at less than 2,500km$^2$. A total of 61 records were retained for analysis. The combined data is reported in Table S1.

For our main analysis we focussed on continental regions as defined by the Natural Earth data at 110m resolution[4]. Of the above records, we manually adjusted the coordinates of 40 observations which were aligned slightly outside the border of our study domain (coordinates reported in columns “Adjusted latitude” and “Adjusted longitude” in Table S1). For each record we then computed the number of HbS alleles and the total number of alleles observed at the location (taking into account HbAA and HbAS samples, and where present HbSS samples) for downstream analysis.

#### 1.3 Component *Pf* global genetic datasets

We collated *P. falciparum* genotype data from the following sources.

##### *1.3.1 MalariaGEN Pf7 and Pf8*

We used previously published data from the MalariaGEN Pf8 resource[5] This includes genotypes from whole-genome sequencing (WGS) from over 24,000 isolates from 97 sites in global malaria-endemic populations. Only samples passing analysis QC filters in that analysis were included in our results; a total of N=24,409 non-excluded samples with *Pfsa* genotype data were included in our analysis. We also use an earlier version of this dataset, MalariGEN Pf7[6], for population genetics analysis as described below.

##### *1.3.2 Molecular inversion probe typing*

We included data from two studies of parasite population genetics based on a molecular inversion probe typing approach, with dense geographic sampling from Democratic Republic of Congo[7] (N=1,436 across 385 locations; additionally a subset of N=392 samples from five locations in Ghana, Tanzania, Uganda and Zambia were also included in this data), and data from multiple regions of Tanzania[8] (N=797 across 13 locations). These samples were typed at a set of over 1,800 SNPs genome wide.

##### *1.3.3 Senegal WGS data*

We included recently published data from multiple regions of Senegal[9] generated using WGS (N=293 across 11 locations).

##### *1.3.4 Uganda molecular surveillance data*

To increase sample coverage we also incorporated previously-unpublished data from Uganda (N=211 across 12 locations). These samples were collected as part of ongoing health facility-based molecular and parasitological surveillance activities. Sample collection has been described previously. For both studies, samples were collected from symptomatic patients aged >6 months after obtaining informed consent. Molecular surveillance samples were collected as dried blood spots (DBS) by finger prick in 16 health facilities across the country between 2016 and 2024. For parasitological surveillance, ~5 mL of blood was drawn into heparin by venipuncture. DBS were made following removal of the buffy coat. Genomic DNA extracted from DBS were subject to two rounds of selective whole genome amplification[10] and libraries prepared using the Watchmaker DNA Library Kit with Fragmentation (Watchmaker Genomics Inc., Boulder, CO), before sequencing using on the Illumina X Plus platform at Psomagen (Rockville, MD). After sequencing, properly trimmed and paired reads were simultaneously mapped onto *P. falciparum* 3D7 (version 3.1) and human genome (version GRCh38) using BWA-MEM. Reads specifically mapping to the *P. falciparum* genome were selected and cleaned using SAMTOOLS and GATK. Samples with high sequencing depth (first quartile of read depth >35X) were retained for variant calling using a *P. falciparum-optimized* GATK4 pipeline[11]. *Pfsa* genotypes were extracted from the filtered WGS variant call format (VCF) file using BCFTOOLS.

##### *1.3.5 Gambia malaria cases and controls (GAMCC) study data*

We also generated new data from samples collected between 1988 and 1989 in Banjul, The Gambia as part of an epidemiological study (GAMCC study; N=345 mild malaria cases). This sample collection has been described previously[12]. In brief, parasite DNA was amplified using selective whole genome amplification[10] and sequencing libraries were prepared using the NEBNext Ultra II FS kit. Samples were sequenced using the Illumina Noveseq 6 platform at the Centre for Human Genetics, University of Oxford. After sequencing, properly trimmed and paired reads were simultaneously mapped onto *P. falciparum* 3D7 (version 3.1) and human genome (version GRCh38) using BWA-MEM. *Pf* genotypes were called using a pipeline based on GATK HaplotypeCaller as previously described[6,13]. Genotypes at *Pfsa* loci were extracted using BCFTOOLS.

#### 1.4 Collation of *Pf* global genetic data

For all studies we extracted genotypes at the lead Pfsa mutations (main text), i.e. those that had the strongest evidence of association with HbS in previous analyses[14,15]. The previously identified lead *Pfsa2* mutation (chr2:814,288 C>T) was not present in the MIP typing data from DRC; in this dataset we instead typed *Pfsa2* using the chr2:814,329 C>T mutation, which is in close linkage and has similar strength of association[13]. For all datasets and loci, we extracted reference and alternate genotypes for the *Pfsa1-4* loci for each sample. For our main analysis we excluded mixed (i.e. heterozygous) genotype calls for each sample at each site (but did not attempt to exclude samples with evidence of high levels of mixture overall).

An interesting feature of the DRC data that distinguishes it from other datasets included here, is that it includes dense geographic sampling with comprehensive coverage of the country, but often with small sample counts at each sampling location despite the overall relatively large sample size. For this reason per-site point estimates of *Pfsa* frequencies in these DRC data are generally highly variable due to low per-site location size, leading to visible variation for these points (small red points in Fig. 1E and Fig. 2). A larger sample from Kinshasa (N=570) is also present in the MalariaGEN dataset.

#### 1.5 Mapping sickle allele frequency worldwide

To create an updated global map of HbS allele frequency, we used a Bayesian geostatistical model. Our approach applies the Integrated Nested Laplace Approximation (INLA), which is a computationally effective model fitting alternative to Markov Chain Monte Carlo and available from the R package R-INLA[16]. In brief, this approach probabilistically models the local frequency of HbS, modelled on the log-odds scale, as following a Gaussian process, that is, its value at any point follows a normal distribution. The Gaussian process is smoothed over space by requiring nearby points to have positive covariance. The survey data are incorporated into this via a binomial likelihood function of the form:

$$P(N_{HbS} = n | N_{total}, f) \approx f^n \cdot (1-f)^{N_{total}-n}, \qquad \text{(S1)}$$

applied to each HbS data record, where the modelled local frequency of HbS at the relevant location is $f$, and the number of observed HbS alleles and total alleles, respectively, are $N_{HbS}$ and $N_{total}$. This approach therefore leads to estimates of the HbS frequency which vary smoothly across geographical space and effectively interpolate between observed data points. To fit the model, we use an implementation in R-INLA, described in further detail in the Supplementary Text. To produce a reasonable global fit, we explored different hyperparameter settings including the range $r_0$ and $\sigma_0$ parameters which control the overall level of smoothness and variance. For our main analysis, we set $r_0$ to 25 and $\sigma_0$ to 0.6, which impose considerable smoothing; additional choices are explored in Fig. S3. From the resulting model fit, for visualisation purposes we then computed the posterior predicted mean, first and third quantiles, and standard deviation of the HbS allele frequency within 5km × 5km grid cells across the study area (Fig. S1).

### 1.6 Aggregating HbS and Pf data across grid cells

To link HbS and *Pf* data across geographical locations, we first discretised the study domain into spatial units. For our main analysis we used regular hexagons with a 1-degree large diagonal length, which corresponds to approximately 110 kilometres at the equator, leading to a total of 6,266 hexagons globally. We also restricted hexagons to land areas by clipping at continent boundaries. A total of 303 hexagonal cells contain *Pf* data survey points. The majority of these cells (260) are within sub-Saharan Africa; 36 and 7 cells respectively lie within Asia and South America. For our main analysis we focus only on the 219 hexagons (198 in Africa) which lie within 200 kilometres of an HbS survey point. These choices are intended to capture the main scope of spatial variation and to reduce any impact of HbS modelling choices on estimation. Results for alternative analyses varying the shape and size of the spatial unit and filtering criteria are presented in Table S3. We found that our results are not substantially affected by these choices, although they lead to minor variation in estimates.

Within each grid cell we aggregated observed *Pf* allele counts and modelled HbS allele frequencies as follows. First, *Pf* allele counts from each study source for the *Pfsa1, Pfsa2, Pfsa3,* and *Pfsa4* variants were summed within each grid cell, forming an overall number of parasites sampled (N) and the number of sickle-associated (*Pfsa+*) alleles observed, separately at each locus. For LD analyses we similarly aggregated combined genotypes across loci in this way (using only samples which had complete genotypes at all loci considered). As described above, mixed (i.e. heterozygous) genotype calls at each locus were excluded from these totals.

To aggregate HbS data while accounting for uncertainty in the HbS map, we first used R-INLA to draw 100 samples from the posterior distribution of the HbS allele frequency map within each hexagon. For each such posterior sample we then computed the median HbS value within the hexagon. In practice, this is implemented as the median value at 50 randomly chosen locations within the hexagon.

This approach therefore produces 100 posterior estimates of HbS allele frequency in each hexagonal cell, drawn from a joint posterior across the global map, along with the aggregated *Pf* survey counts by source study. The analysis described in main text uses these values to consider the uncertainty in HbS and *Pfsa* allele frequencies.

### 1.7 Comparison to Piel et al HbS map

To compare our HbS allele frequency estimates with those of Piel et al.[3], we further aggregated the Piel et al map across hexagons and plotted separately across continents (Fig. S2). Overall, our results are in line with Piel ($R = 0.94$, P-value $< 0.001$).

### 1.8 Modelling the relationship between HbAS/SS and *Pfsa*+ genotypes across geographical areas

To model the effect of HbS on *Pfsa+* prevalence at hexagonal level in the study domain, we used a regression approach that accounts for HbS allele frequency as the predictor and *Pf* genotype counts as the outcome variable. The model is based on the Besag-York-Mollié[17] (BYM) framework with adaptations to our setting as follows. *Pfsa* genotype counts are modelled through a binomial likelihood function, which accounts for variation due to the finite sample size at each *Pf* data point. The *Pfsa+* frequency parameter $f_j$ in grid cell $j$ is then modelled on a transformed scale according to a regression link function. Although we considered various possible link functions; the analysis described in main text uses a generalised logistic link (denoted gl()) which provides a relatively flexible S-shaped model of the relationship between HbS and *Pfsa+*. Specifically, we used the following form:

$$f = \mathrm{gl}(\pi) = \frac{1}{(1 + e^{-\pi})^{\frac{1}{\nu}}}. \qquad \text{(S2)}$$

In this formula, $\pi$ is the transformed frequency and $\nu > 0$ is an additional parameter which controls the form of the relationship. Similar to the regular logistic function (which is obtained when $\nu = 1$) the generalised logistic function represents an S-shaped curve which asymptotes, respectively, to 0 and 1 for small and large values of $\pi$. In regular logistic regression, $\pi$ is interpreted as the log-odds corresponding to frequency $f$, but in the more general case $\pi$ is the log-odds corresponding to $f^{\nu}$, i.e. to the frequency raised to the $\nu$th power.

For a regression framework we then model the transformed frequency $\pi = \pi_j$ in each cell j in terms of the estimated HbS frequency. Specifically, we assume that $\pi$ takes the form:

$$\pi = \beta_0 + \beta_1 f_{HbAS/SS} + \zeta \qquad \text{(S3)}$$

Here, $\beta_0$ and $\beta_1$ are parameters that control the overall *Pf* frequency and the strength of the relationship between HbS and *Pf* on the transformed scale, and $\zeta = \zeta_j$ is a quantity that accounts for possible additional variation in cell j that might vary either with geographical location or with sampled points. In our analysis, we assume $\zeta$ is a sum of two independent variables, i.e. $\zeta = \omega + \eta$, where ω is represents a geographically structured effect that induces correlation between neighbouring grid cells (i.e. those that share a boundary segment) and η accounts for additional independent (non-geographical) variation in the value of $\pi$ for each sample. A rationale for this formulation is that, regardless of any potential effect of HbS, *Pf* allele frequencies are the result of population genetic processes including genetic drift and migration that are likely *a priori* to generate spatial autocorrelation; at the same time each *Pf* datapoint arises from specific sampling schemes and at various time points in each area which may also generate sample-specific variation.

In the approach taken here, both ω and η are modelled as gaussian random effects and we follow the BYM2 formulation[18] in writing their variances as:

$$\text{var}(\omega) = \sigma^2\phi \text{ and } \text{var}(\eta) = \sigma^2(1-\phi) \qquad \text{(S4)}$$

where $\sigma^2 > 0$ represents the overall scale of the random effect. The combined variance of the random effect is therefore $\sigma^2$, and $0 < \phi < 1$ determines the proportion of total variance attributable to spatial dependence.

Given *Pfsa* allele counts and values of HbS allele frequency in a set of grid cells, we fit the above model using Template Model Builder (TMB)[19]. In the main analysis we used the following prior specification which we implemented as penalty terms in the marginal log-likelihood function:

- $\beta_0$: very weak Gaussian $N(0, 100^2)$ prior
- $\beta_1$: weakly informative / regularising Gaussian $N(0, 10^2)$ prior.
- $\nu$: weakly informative / regularising Gaussian $N(0,1)$ prior on log $(\nu)$.
- $\phi$: weakly informative Gaussian $N(0,10^2)$ prior on log-odds $(\phi)$.

Finally, for the random effect standard deviation parameter $\sigma$ we follow the PC prior specification[18]. Specifically, we assume that it is relatively unlikely that the standard deviation $(\sigma)$ of the random effect is larger than 1 (i.e. that this occurs *a priori* with 1% probability), leading to an exponential prior on $\sigma$ with rate $\log(0.01)$[18]. In practice this is implemented using a Type II Gumbel distribution on $\tau = 1/\sigma$ as described in Section 4.1 of Riebler et al[18]. The prior choices given above are intended to be largely uninformative - that is, to provide minimal influence on the model fit—while still regularising estimates in cases where little information is present in the data. In our analysis, this would occur for example if the analysis were applied to regions where there is little variation in the HbS frequency.

TMB uses a Laplace approximation approach to integrate out the random effects from the marginal loglikelihood[19]. Model fitting returns both the posterior mode and a covariance matrix of the fixed parameters (i.e. $\beta_0, \beta_1$ and, for the generalised logistic approach, log $(\nu)$). This therefore provides a gaussian approximation to the posterior and we exploit this to sample $n = 100$ parameter sets from this approximate posterior distribution, for each model fit. In our analysis, the HbS map is itself the result of a statistical model and, as described above, we have represented this uncertainty with posterior samples. To handle this, we re-fit the above model 100 times based on 100 draws from the joint posterior distribution of HbS maps. In total this can therefore be considered to generate $n = 10{,}000$ draws from the joint posterior of HbS and *Pf* data.

#### 1.9 Global and regional analysis, and model sensitivity

We fit the above model using either all global data or using only data from specific regions including all Africa (Fig. 1C,D), sub-regions of Africa (Fig. 2), and individual countries or groups of countries where there was sufficient geographical spread of data points to make the estimates (Fig. S4). For analysis in specific areas, we first defined regions as a list of relevant countries and clipped hexagons at these region boundaries. We then repeated the analysis using only Pf data points (and HbS allele frequency estimates) within the clipped hexagon boundaries. We note that analysis in smaller regions necessarily involves fewer data points, and many of our estimates in individual countries have large credible intervals (Fig. S4 and Table S3).

Since the model above also relies on discretising space into hexagons, we also tested robustness to different specifications including using hexagons with either 1°, 1.35°, and 2° long diagonal, or square cells with 1°, 1.35°, and 2° edge length. We found minimal differences in analysis results using these specifications (Table S3). For our main analysis we use relatively small hexagonal cells to best capture the variation in host and parasite allele frequencies at local scales.

#### 1.10 Summarising the joint HbS-*Pfsa* model fit

To visualise the joint HbS-*Pfsa* model fit (Fig. 1C, D and Fig. 2A to 2D), we plot the mean HbS estimate (averaged across posterior draws) against the point estimate of frequency from the *Pf* data in each hexagon. We then overlayed lines/areas representing the mean and the 2.5% and 97.5% quantiles of the *Pfsa* allele frequency predicted by the *Pfsa*+-HbS regression model across values of $f_{\text{HbAS/SS}}$.

In the regression framework above, the estimated $\beta_1$ parameter represents the strength of association. However, $\beta_1$ is difficult to interpret directly because it relates to the scale of the linear predictor which, as described above, is on a generalised log-odds scale which is dependent on the parameter $\nu$. We therefore instead focussed on a more interpretable 'slope' of the model, which we define as the difference in predicted *Pfsa*+ frequency at $f_{\text{HbAS/SS}} = 10\%$ and $f_{\text{HbAS/SS}} = 20\%$. This quantity, which we denote by e.g. $\Delta f_{1+}$ for *Pfsa1*+, can be computed for each of the 10,000 posterior draws from the joint posterior of the model to identify its posterior mean and 95% credible interval. These values are reported in main text, Fig. 2B and Table S2.

#### 1.11 Between-locus linkage disequilibrium is not associated with local HbS frequency

A surprising feature of the *Pfsa*+ alleles is that they are in strong linkage disequilibrium (LD) - that is, they tend to be observed together in the same infection – despite the fact they lie in separate regions of the *Pf* genome. This is notable because meiosis is expected to rapidly reduce LD between variants on different chromosomes – or distal on the same chromosome – provided there is at least modest levels of outcrossing.

Analysis across geographic areas in our data demonstrates that LD between *Pfsa1*+ and *Pfsa3*+ is positive – that is, parasites with both alleles are more common than would be expected if the loci were statistically independent – everywhere in Africa (fig S8A). In many locations the LD is very high but a smaller number of sites do show lower values, notably including areas of Gambia, Gabon, and Benin and to a lesser extent Tanzania and Cameroon. Analysis of joint genotypes at the two loci shows that this is generally caused by an increased frequency of *Pfsa1*- / *Pfsa3*+ parasites, which in some regions reach frequencies over 20%, as is consistent with the higher frequency of the *Pfsa3*+ allele (Fig. S8B). However, we did not observe any strong relationship between HbS frequencies and parasite LD (Fig. S8C). Indeed, high LD (D' > 0.6) is observed across geographical areas and across the range of observed $f_{\text{HbAS/SS}}$ values, while lower-LD areas are also widely distributed. Similar conclusions can be drawn when considering all three loci prevalent in east or west African regions (i.e., *Pfsa1*, *Pfsa3*, and *Pfsa4* in west Africa; *Pfsa1*, *Pfsa2*, and *Pfsa3* in east Africa; Fig. S8, D and E).

The reasons for the elevated LD are not currently known, but possible explanations include biological interactions between the loci – i.e. functional epistasis – or, as we explore in main text, potential maintenance of LD through co-selection of the loci across geographical areas.

#### 1.12 Theoretical implications of the geographical comparison for parasite relative fitnesses

##### *1.12.1 Single-locus model in a homogeneous population*

The relationship between HbS genotype and *Pfsa*+ frequencies shown in Figs. 1-2 suggests that the *Pfsa*+ alleles may be positively selected (or at least not negatively selected) in host populations where HbS is highly prevalent, and vice versa in populations where HbS is at low frequency. To provide some information about the underlying parasite fitnesses, we considered the implications of this observation under a simplified model of epidemiological fitness. To describe this, we focus on a single *Pfsa* locus and express results in terms of the fitness of a parasite at the time of infection, given its genotype and that of the host. In the following we denote this fitness by $\gamma_{ph}$ where $p$ and $h$ are the parasite and host genotypes respectively. Thus, fitnesses form a matrix:

$$\Gamma = \begin{pmatrix} & AA & AS/SS \\ - & \gamma_{-A} = 1 & \gamma_{-S} \\ + & \gamma_{+A} & \gamma_{+S} \end{pmatrix}. \qquad \text{(S5)}$$

For concreteness, 'fitness' here refers to fitness at time of infection, and specifically to the expected number of transmitted offspring that a parasite with a given genotype has when infecting a host with the given genotype. As shown above, we consider all fitnesses as relative to the fitness $\gamma_{-A} = 1$ of wild-type (*Pfsa*-) parasites infecting AA individuals, and we group individuals with HbS genotypes for this discussion (S = HbAS or SS). The use of relative rather than absolute fitnesses means that the analysis focusses on the relative frequencies of the two alleles (in a parasite population which is assumed to remain sizeable) and therefore does not address small-population effects such as genetic drift.

With this framework a simple model of parasite allele frequency evolution can now be set up by imagining a mosquito biting at random in large host population with a frequency $f_{\text{HbAS/SS}}$ of AS/SS genotypes and $f_{\text{HbAA}} = 1 - f_{HbAS/SS}$ of AA genotypes. If the frequency of + and – parasites in the biting mosquito population is $f_+$ and $f_- = 1 - f_+$, this leads to frequencies in the host population after biting of

$$f_+^{new} = \frac{f_+(f_A\gamma_{+A} + f_S\gamma_{+S})}{f_+(f_A\gamma_{+A} + f_S\gamma_{+S}) + f_-(f_A\gamma_{-A} + f_S\gamma_{-S})} \,. \tag{S6}$$

This formula is derived by computing the total number of individuals successfully infected with *Pfsa*+ parasites and the total number infected by any parasite; the ratio of these is (S7). This formula therefore represents a highly simplified model of parasite evolution which, however, captures the effects of parasite fitnesses dependent on host and parasite genotypes.

Iterating (S7) shows that in a homogeneous population (i.e. where mosquitoes bite at random), except for a specific value of $f_S$ dependent on the fitnesses, $f_+$ will either rapidly increases to fixation, or decreases so that the + allele is lost. Specifically, positive selection occurs whenever

$$f_{\text{HbAS/SS}}(\gamma_{+S} - \gamma_{-S}) + f_{\text{HbAA}}(\gamma_{+A} - \gamma_{-A}) > 0 \,, \tag{S7}$$

and negative selection if this quantity is less than zero. Only relative fitnesses are important in this equation, so as above we may take $\gamma_{-A} = 1$. For a fixed value of $\gamma_{-S}$ positive selection occurs if

$$\gamma_{+S} > -\frac{f_{\text{HbAA}}}{f_{\text{HbAS/SS}}}\gamma_{+A} + \frac{f_{\text{HbAA}}}{f_{\text{HbAS/SS}}} + \gamma_{-S},$$

and negative selection if the inequality is reversed. Although this conclusion only applies to a homogenous population (i.e. where mosquitoes bite at random) and is no longer true if there is heterogeneity e.g. in the A and S allele frequencies (as described below and in Fig. 3C). However, it is instructive to consider the implications as if population were homogenous. Applying this to the data in Fig 1C suggests that – all else being equal – there is likely no negative selection against *Pfsa*+ in the right-hand part of the graph, e.g. when $f_{HbAS/SS} \geq 25\%$, and likely no positive selection for *Pfsa*+ in the left-hand side of the graph, e.g. when $f_{HbAS/SS} \leq 5\%$. Lines $l_1$ and $l_2$ of Fig. 1A show the resulting constraints under the assumption that HbS genotypes strongly impede infection by *Pfsa*- parasites ($\gamma_{-S} = \frac{1}{100}$). Similar results are found under higher choices of this parameter (Fig. S5).

We note that the parameters considered here pertain to the overall fitness of parasites rather than the effect on disease outcomes. As such, no direct estimates of these parameters are available from current epidemiology studies. However, various estimates of the protective effect of HbS on severe and milder forms of infection have been made[20] – without taking into account parasite genotype – and broadly, these estimates suggest that HbS genotypes have a strong protective effects against severe disease (e.g. $RR = \frac{1}{5}$ to $\frac{1}{10}$), weaker effects against asymptomatic disease, and intermediate protection against mild disease. Accordingly, line $l_3$ in Fig. 3A corresponds to region below which HbS genotypes confer at least some protection against the overall population of parasites, even when *Pfsa*+ parasites are at high frequency ($f_+ = 50\%$).

Two notable features of Fig 3A are that the bounded region suggests the fitness cost of *Pfsa*+ parasites in A individuals ($\gamma_{+A}$, x axis) must be less than 100%, but it is nevertheless substantially larger than $\gamma_{-S}$. That is, *Pfsa*+ alleles are less fit in HbAA individuals, but this effect is much weaker than the protective effect of HbS on *Pfsa*- parasites. Secondly, it is notable that the possible strength of effect of HbS against *Pfsa*+ parasites ($\gamma_{+S}/\gamma_{-S}$, y axis) could apparently take a range of values, including situations where there is no protective effect at all ($\gamma_{+S} = \gamma_{+A}$) or even, potentially, risk effects.

These parameters can also be compared to the previous estimates of relative risk of severe malaria using severe infections from The Gambia and Kenya[13]. Roughly speaking these suggest that protection due to HbS is very strong (on the order of $RR = \frac{1}{100}$) for *Pfsa*- parasites, and potentially close to no protection against parasites carrying all the *Pfsa*+ alleles. Thus, these estimates (although not direct estimates of the fitnesses considered here) nevertheless appear compatible with the above analysis of population frequencies.

#### 1.12.2 *Multi-locus model in a geographically heterogeneous population*

To provide a more realistic analysis we extended the above model to capture spatial heterogeneity and to a multi-locus system. A simulation using this model is shown in Fig. 3C-E. Full details are given in Supplementary Text, but the two important features are as follows.

First, the model is extended to allow for geographical heterogeneity in the frequency of HbAS/SS genotypes. Conceptually, we achieve this by modelling parasite populations separately at different locations and by assuming mosquitoes bite locally (i.e. that infections are sourced from nearby cells, according to some biting distance distribution which must be chosen). The fitness of different infections is then computed using a modified form of (S7) based on the local frequency of HbAS/SS genotypes. Importantly, as discussed in main text, this alters the behaviour of the evolution by (S7) such that (for some parameter values, including those in the form discussed above) the allele frequencies reach a stable equilibrium.

Second, we also extended the model to account for multiple loci – specifically, to two loci in our implementation. (Conceptually we think of these as potentially corresponding *Pfsa1* and *Pfsa3*.) This entails extending the matrix (S5) to account for the fitness of genotypes at two loci, i.e. to a matrix of the form

$$\Gamma = \begin{pmatrix} & AA & AS/SS \\ -\,- & \gamma_{--A} = 1 & \gamma_{--S} \\ -\,+ & \gamma_{-+A} & \gamma_{-+S} \\ +\,- & \gamma_{+-A} & \gamma_{+-S} \\ +\,+ & \gamma_{++A} & \gamma_{++S} \end{pmatrix}. \qquad \text{(S7b)}$$

The rate at which these genotypes is produced is moreover controlled by allowing a proportion ($m$) of mosquitoes to bite twice, after which meiosis occurs (the two loci are assumed to lie on different chromosomes). This model is therefore substantially more complicated than the single-locus model (or equivalently the model with no outcrossing) as the ultimate evolution depends both on the starting frequencies, the fitnesses, and the rate of outcrossing. Nevertheless, we found that similar results to the single-locus model are obtained with reasonable choices of fitness in (S7b).

Given the high level of LD observed between *Pfsa* loci, a question of fundamental interest is whether they are involved in epistatic interactions. (This could in principle occur within AA individuals, or within HbS-carrying individuals, or both). In the context of our model, this might be said to occur if the fitnesses for +- and -+ genotypes deviate from simple models that place them intermediate to those for the – and ++ genotypes. As shown in Fig. 3E, we found that substantial LD can be obtained from models with no interaction (e.g. additive or multiplicative models). However the parameter choices that best fit observed frequency data are not of this form and do suggest a form of epistasis between the loci.

A full description of the model is given in Supplementary Text. To implement the model we used the WebGPU framework within a webpage implementation, which allows interactive exploration of the parameter space. The simulation can be found at https://www.chg.ox.ac.uk/~gav/projects/pfsa/simulation/.

### 1.13 Curation of MalariaGEN Pf7 population-genetics dataset

For haplotype and selection analyses, we used genome-wide genotype calls from the MalariaGEN Pf7 resource[6]. We post-processed these data with the aim of generating a robust set of genotype calls at bi-allelic variants across the genome, suitable for population genetic analyses as follows.

#### 1.13.1 *Data sources and preprocessing*

Data was downloaded from the MalariaGEN Pf7 data release web page (https://www.malariagen.net/resource/34/); see also https://apps.malariagen.net/apps/pf7/ for more information. We downloaded genotype data for all samples encoded in VCF format files. To make these easily processable, we undertook an initial round of filtering which involved removing all data fields other than genotypes (GT), removing all alleles with low frequency (minor allele count < 20) across the whole dataset. We noted that some genotypes in the data are called in 'phased' form (e.g. as GT=0|1) whilst most are unphased (GT=0/1); this is a feature of the GATK output. Because this prevented some downstream tools from working, we replaced these genotypes with unphased genotypes using a GNU sed command.

#### 1.13.2 *Dataset filtering and phasing*

We restricted attention to samples from African populations passing Pf7 QC filters (N=8492) and further restricted to largely unmixed infections (determined as having Fws>0.9[21]) leaving N=4,788 samples for analysis. We also restricted attention to PASS variants (which excludes variants outside the core genome). The Pf7 dataset contains many variants called as multiallelic, but where the third or additional alleles are at very low frequency. We used vcflib's vcffilter to remove alleles with < 20 alternate allele calls from multi-allelic variants, treating the remaining variants as biallelic if only two alleles remained. We excluded remaining multi-allelic variants from the dataset.

To produce a fully phased dataset with no missingness in this set of largely unmixed samples, we then set all heterozygous calls to missing and re-phased the data (which fills in missing genotypes) using BEAGLE v5.4 with 24 phasing iterations. Due to the algorithm stochasticity, a small number of heterozygous genotypes remained in the output files and these were assumed to be homozygous for the reference allele. The output of this step contained a total of N = 2,959,655 bi-allelic variants phased with no mixed or missing genotypes.

#### *1.13.3 Determination of ancestral and derived alleles in the Pf genome*

Most population-genetic analyses require alleles to be polarised (in the sense that the ancestral and derived alleles have been identified). We therefore generated a set of ancestral/derived allele calls using the program est-sfs (version 2.04)[22]. First, a multiple sequence alignment of the *P. falciparum* 3D7 reference sequence, an assembly for the related Gorilla malaria parasite *P. praefalciparum* (PfG01), and an isolate of the chimpanzee infecting *P. reichenowi* (PrG01) was computed using the software MAFFT for each chromosome (using the –auto command line argument and four threads). We then extracted the nucleotide base at the positions of all variants in the MalariaGEN Pf6 resource which were marked as 'PASS', and input these into est-sfs. Data was formatted such that *P. praefalciparum* was considered the first outgroup (i.e. the closer relative to *Pf*) and *P. reichenowi* a more distantly related outgroup[23]. Based on this, ancestral state probabilities were determined using the maximum likelihood method and the Kimura 2-parameter model. This models the rate of transitions (substitutions between pyrimidines or between purines) and the rate of transversions (purine to pyrimidine and vice versa) across the genome in order to estimate genetic distances. The nucleotide type with the highest probability was determined to be the ancestral state allele.

We used QCTOOL (https://www.chg.ox.ac.uk/~gav/qctool) with this file to polarise the phased genetics dataset for downstream analyses. We refer to this as the popgen dataset below.

#### *1.13.4 Computation of selection metrics*

We used selscan to compute metrics of positive selection (iHS and iHH12) across all chromosomes within each country, and either across all years or within specific year groups (before 2000, 2000-2009, 2010-2019, and 2020s).

We also used a custom R script to implemented a version of the Beta[24,25], Dango[26], and Tajima's D statistics. These statistics were computed in 5kb or 10kb windows centred on each variant with at least 1% allele frequency in each population. For Beta, we set the *p* parameter (which controls how strongly local variation affects the statistic) to 20 or 50. At each focal variant we also computed the total nucleotide diversity (i.e. the average number of mutations separating samples within the window), and the proportion of nucleotide diversity that was due to differences between the two alleles (i.e. between samples exactly one of which carries the derived mutation at the focal variant).

#### *1.13.5 Genealogy estimation*

To produce more accurate genealogical estimates, we used the program RELATE[27] to estimate an ancestral recombination graph across the genome. We used initial parameters of Ne=100,000, and a constant recombination rate of 1cM ~ 17kb. Mutation parameters were set following the calculation by Otto et al[23]. Specifically, we assumed $3.83 \times 10^{-10}$ mutations and four mitoses per erythrocyte cycle, and between 66 and 336 erythrocyte cycles per generation (i.e. between consecutive meioses), leading to per-generation mutation rates of between $6.3162 \times 10^{-9}$ and $3.21552 \times 10^{-8}$. Following initial estimation, we re-estimated the population size, coalescence rates, and branch lengths using the RELATE 'EstimatePopulationSize' script and extracted the estimated genealogical tree for *Pfsa1* at chr2:631,190. For visualisation purposes, we then extracted the tree at the *Pfsa* loci in newick format.

#### *1.13.6 Haplotype visualisation*

To illustrate local haplotypes patterns, we plotted local haplotypes alongside an indicator of population origin using the ordering compatible with the estimated genealogy (Fig. 4 and Fig. S8). For visualisation purposes, the datasets were first downsampled by grouping by country and *Pfsa*+ genotype and then randomly sampling at most 25 samples in each group for visualisation. We annotated variants putatively linked to the lead *Pfsa*+ mutation using the following criteria:

- Within 5kb of the lead mutation

- Frequency at least 2% across the whole sample
- Frequency < 5% on Pfsa- background
- Frequency at least 5 times higher on *Pfsa*+ than *Pfsa*- background

In Fig. 4 legend, for simplicity we report simpler criteria (at least 10 occurrences on *Pfsa*+ background and at most 10 on *Pfsa*- background) - but we verified that these criteria produce the same variants as above. For tree annotation, we assigned mutations to the most ancestral branch such that 99% of descendant tips carry the mutation.

Due to the use of two different mutation rate estimates (per transmission cycle) we annotated the genealogy with time interpreted as number of transmission cycles in the past, expressed as a range based on the two possible values. Specifically we used the trees estimated using the smaller mutation rate estimate ($6.3162 \times 10^{-9}$), i.e. fewer mutations per transmission, and scaled values in the legend to also annotate the corresponding times based on the larger mutation rate.

The trees output by RELATE include branch lengths; additionally, it is possible to estimate the posterior distribution of allele ages by sampling branch lengths using an MCMC method[27]. After tree estimation, RELATE works with a single tree topology at each site, so this method does not resample tree topology. To gain age estimates, we first identified the branch corresponding to the relevant *Pfsa*+ mutation as above, and then computed the age of the upper and lower end of the branch across each of N=100 MCMC-sampled branch length estimates. The 95% and 80% quantiles of these upper and lower estimates are shown below the tree in Fig. 4 and Fig. S8.

##### *1.13.7 GEVA-style allele age estimate*

As an alternative way to estimate allele age, we adopted a method based on counting the number of mutational differences between pairs of samples in a region around the *Pfsa1*+ mutation. This is motivated by the method in GEVA[28], but we based analysis on mutations in a fixed 5kb region around the lead *Pfsa1*+ mutation and only considered a mutational clock (i.e. we did not try to take into account possible information from inferred recombinations). One mutation, at position chr2:630,737 C>T, was at intermediate frequency on both *Pfsa*+ and *Pfsa*- backgrounds and therefore does not fit with the tree topology. Inspection of the raw Pf7 data indicates this variant overlaps a nearby insertion/deletion. We removed this variant before computation.

### Supplementary Text

#### 1.14 Estimation of global HbS frequency map using R-INLA

We use R-INLA[16] to estimate and map the HbS frequency surface. This approach uses a specific form of covariance function between locations (the Matérn correlation function, described below) which for values $v_X, v_Y$ at two locations $X, Y$ takes the following form:

$$\text{Matérn covariance}\,(\mathrm{v_X}, v_\mathrm{Y}) = \frac{\sigma_\zeta^2}{\Gamma(1)}(\kappa \cdot \delta_{XY})^1 K_1 \kappa\, \delta_{\mathrm{XY}}\ . \tag{S21}$$

In this formula, $\delta_{XY}$ denotes the geographic distance between locations X and Y, while $\kappa$ is a parameter that controls the smoothing and $\sigma_\zeta^2$ is the overall variance; also $K_1$ denotes the modified Bessel function of the second kind and $\Gamma(1)$ the Gamma function evaluated at the value 1. This function equals $\sigma_\zeta^2$ when $X = Y$ and smoothly decays to close zero beyond a distance of around $r = \sqrt{8}/\kappa$[29]. In the following we refer to $r$ as the range parameter.

A full description of the model is then specified as follows:

$$\begin{gathered} N_{HbS} | N_{total}, f_X \sim \text{binomial}(f_X, \mathrm{N}_{total}) \\ \text{logit}(f_X) = \log\left(\frac{f_X}{1 - f_X}\right) = \mu + \zeta_X \\ \theta \sim \text{prior}(\theta) \end{gathered} \tag{S22}$$

Where $N_{HbS}$ and $N_{total}$ are the number of HbS alleles and the total number of alleles in a data record at location $X$, $f_X$ denotes the modelled frequency of the HbS allele at location $X$, and $\theta = (\mu, \sigma_\zeta, r)$ are model parameters reflecting the mean allele frequency and the variance and range of the spatial effects respectively. In (S22), the linear predictor includes the global mean $\mu$ and a random effect $\zeta_X$, which is assumed to follow a zero-mean

Gaussian distribution following the Matérn covariance framework outlined above, and therefore links the modelled frequency between nearby locations. The parameters of this covariance function ($\sigma_\zeta$ and the range parameter $r$) and the fixed effect $\beta_0$ can be learnt from the data, but in practice for our analysis we chose to fix them as detailed below.

The use of the Matérn covariance function is motivated by previous work[29] which demonstrated that processes defined in this way can be effectively approximated by working over a triangular mesh. Accordingly, we use R-INLA functionality to define a global mesh (with vertices at observed survey points) and to estimate the model parameters. The model is fit under a Bayesian framework but for our analysis we used an uninformative prior on the global mean (uniform between $-\infty$ and $\infty$), and used specific choices of Matérn covariance parameters to give reasonable levels of smoothing. Specifically, for the Matérn parameters, we used a simple manual approach (guided by visualisation of the resulting maps) to choose variance and range parameters; for our main analysis we set $\sigma_\zeta$ to 0.6 and $r$ to 25°, corresponding to about 2,750 kilometres at the equator. (We note that R-INLA also internally adds a small i.i.d. random effect term with fixed precision to S23 to provide model robustness[16]). These prior choices lead to an estimated HbS frequency surface that is relatively well smoothed out over large distances and where the variability away from data points is limited (Fig. 1). The above approach allows us to obtain posterior estimates of the global HbS frequency surface (e.g. the posterior mean, depicted in Fig. S1) and enables posterior samples to be drawn which we use in downstream analysis. Sensitivity of our analysis to these choices is further explored in Fig. S2. The model predictions at any geographical location, including rectangular pixels for plotting or *Pf* survey points, can then be extracted from the approximation.

### 1.15 Simulation of parasite allele frequency evolution

To investigate parasite evolution we developed a discrete-time, deterministic simulation of genotype frequencies. This is described in the following sections.

#### *1.15.1 Simulation scenario and notation*

We consider a geographical space divided uniformly into small grid cells. Each cell $z$ is assumed to contain a host population made of up individuals who carry the HbS allele (with frequency $f_S(z)$) or else the HbAA genotype (with frequency $f_A(z) = 1 - f_S(z)$). These frequencies vary with geography, and in our simulation we use the estimated posterior mean frequencies from our HbS map (Fig. 1B).

We also consider infecting parasites having genotypes at two biallelic loci. Genotypes at these two loci are encoded as $g_{Pf} \in \{--, -+, +-, ++\}$, where e.g. $g_{Pf} = -+$ means that the parasites have the $-$ allele at the first locus and the $+$ allele at the second locus. In the main simulation, these are arranged so that the $+$ allele is associated with increased ability to infect S genotype individuals, relative to the $-$ allele. For concreteness we note that all parasites are haploid (except that they combine for meiosis during transmission, as described below).

#### *1.15.2 Infection model*

At each timepoint *t*, in each cell, a subset of host individuals are infected by parasites which are sampled in nearby cells (based on frequencies at time t-1) by mosquitoes which then fly in and bite individuals in the local population. Infections succeed or fail based on the host and parasite genotypes, and the successful infections form the new local parasite population at timepoint *t*. Specifically, to model host-parasite interaction, we assume that the probability that a bite by an infected mosquito leads to a successful infection (i.e. one that contributes to the new local population) is given by a fitness matrix Γ which is of the following form:

$$\Gamma = \begin{bmatrix} & A & S \\ -- & 1 & \gamma_{--,S} \\ -+ & \gamma_{-+,A} & \gamma_{-+,S} \\ +- & \gamma_{+-,A} & \gamma_{+-,S} \\ ++ & \gamma_{++,A} & \gamma_{++,S} \end{bmatrix}. \quad \text{(S8)}$$

Here $\gamma_{g,h}$ denotes the relative probability that an infection of a host with genotype h by a parasite with genotype g will be successful. As shown, all fitnesses are interpreted relative to that of wild-type ($--$) parasites infecting A individuals, and this fitness is therefore taken as $\gamma_{--,A} \equiv 1$. We discuss choices of these parameters commensurate with current data further below, but to fix ideas we typically consider fitness effects of the following form:

$$\Gamma = \begin{bmatrix} & A & S \\ -- & 1 & \ll 1 \\ \mp & \text{intermediate} & \text{intermediate} \\ \pm & \text{intermediate} & \text{intermediate} \\ ++ & < 1 & < 1 \end{bmatrix}. \quad \text{(S9)}$$

These choices mean that the + alleles at both loci are associated with strongly increased fitness in S individuals (relative to the $--$ genotype), but with modestly decreased fitness in A individuals.

*1.15.3 Transmission model.*

Denote the frequency of the four genotypes in a given cell z as $f_{--}(z)$, $f_{-+}(z)$, $f_{+-}(z)$, and $f_{++}(z)$. If we wish to emphasise the timepoint we will add a superscript t. We will also denote host genotypes as $h \in \{A, S\}$ and parasite genotypes as $g \in \{--, -+, +-, ++\}$.

Consider a mosquito which draws a parasite-infected bloodmeal in cell $w$ at timepoint $t$, and then bites a host at target cell z. Let $\Omega_{h,g} = \Omega_{h,g}(w, z)$ be the combined probability that this leads to a successful infection of a host of genotype h with a parasite of genotype $g$. This is equal to the probability the mosquito sampled a parasite of genotype $g$, times the probability of infecting an individual of genotype $h$, times the infection fitness:

$$\Omega_{h,g} = f_g^{t-1}(w) \cdot f_h(z) \cdot \gamma_g \,. \quad \text{(S10)}$$

The above describes mosquitoes which sample and transmit a single parasite. To study breakdown of linkage disequilibrium patterns, we further extended the model to allow for meiosis. Briefly, we assume that a proportion $\theta$ of mosquitoes bite twice (sampling two parasites at random from the source cell w). These undergo meiosis and produce four offspring gametes each of which is transmitted with probability $\frac{1}{4}$. In our simulation, we assume that the two loci are on different chromosomes so that they are transmitted to daughter gametes independently. The remaining mosquitoes sample only a single parasite. In symbols, let $M_g = M_g(w)$ denote the probability that the transmitted offspring from a mosquito biting in cell w has genotype g. This can be computed by summing over the possible parasite genotypes that the mosquito may draw up in a bloodmeal, i.e. as:

$$M_g = \sum_{g_1} \sum_{g_2} f_{g_1}(w) f_{g_2}(w)\, P(\textit{transmitted offspring of}\, g_1, g_2 = g)\,, \quad \text{(S11)}$$

then the full expression for $\Omega_{h,g}$ becomes:

$$\Omega_{h,g}(w, z) = \left[(1-\theta) \cdot f_g^{t-1}(w) + \theta M_g(w)\right] \cdot f_h(z) \cdot \gamma_g \quad \text{(S12)}$$

Equation (S12) therefore represents a simplified model of transmission of parasites between cells, taking place via mosquitoes that sample parasites in cell w and transmit them to cell z, and accounting for the effects of meiosis in shuffling genetic material in a proportion of transmissions.

*1.15.4 Evolution update model.*

Formulae for the updated allele frequencies at cell z can now be worked out by counting successful infections with each genotype. For concreteness, suppose there are $N = N(z)$ individuals in cell z and that each individual is bitten with probability $\lambda = \lambda(z, w)$ by an infected mosquito flying from cell w. Then according to the above, the expected number of successful infections in cell z originating in cell w and with the given host genotype h and parasite genotype g is $N\lambda\Omega_{h,g}$. The total number of successful infections originating from all source cells with the given parasite genotype g is therefore obtained by summing over cells and host genotypes:

$$T_g^t(z) = N \cdot \sum_w \lambda(w, z)\big(\Omega_{A,g}(w, z) + \Omega_{S,g}(w, z)\big) \quad \text{(S13)}$$

Assuming that biting rates are small enough that each host is infected at most once, the total number of infections at timepoint t in cell z and the updated allele frequency of genotype g can then be worked out as:

$$N^t(z) = \sum_{g'} T^t_{g'}(z) \;\; g' \in \{--, -+, +-, ++\}$$

and (S14)

$$f^t_g(z) = \frac{T^t_g(z)}{N^t(z)}$$

Formula (S14) is the counterpart of (S6) in this spatially variable setting. It can be regarded as reflecting the new expected frequency of parasite genotype $g$ at time $t$ in cell $z$; or alternatively as approximately holding in a population where both the number of individuals N(z) and the number of infecting mosquito bites is sufficiently large that stochastic effects can be disregarded.

We note that (S14) is similar to models that have been extensively studied in the literature. For example, it is similar to formulae (10a-11d) in Lewontin and Kojima[30] who studied linkage disequilibrium in a homogeneous diploid population, in particular under heterozygote advantage models; and to formula (1.2) in Karlin[31] who studied conditions for 'protected' polymorphism, i.e. the stability of polymorphic equilibria, at a single locus. Nei and Li[32,33] demonstrated that stable patterns of linkage disequilibrium between pairs of loci arise in populations with spatially varying patterns of selection, even when there is no epistasis between loci. More detailed mathematical results are also available[34-38], detailed in Bürger[35], including formal results on asymptotic behaviour in limiting cases such as where the migration patterns are weak (i.e. the population is close to a set of independent populations) and/or strong (i.e. the population is similar to a single panmictic one).

#### *1.15.5 Simulation transmission weighting model.*

The above model includes terms $\lambda(w, z)$ defined the probability that an individual in cell z is bitten by a mosquito from cell w at each timepoint. These can therefore be interpreted as weights that affect the contribution of cell w to z on each iteration. Since an important feature of the model is that parasites migrate locally, we modelled these using a distribution biting distance which decays to zero beyond a radius around each cell. Specifically, for a flexible form we assume a maximum distance D, and assumed that the distances flown by mosquitoes is governed by a Beta distribution as:

$$\lambda(w, z) = \mathrm{Beta}_{1,r}\left(\frac{\mathrm{distance}(z, w)}{D}\right) \quad \text{(S15)}$$

where $r$ is a shape parameter that controls the shape of the beta function. This form was chosen for flexibility; the specific form of this biting distribution is relatively unimportant provided bites are concentrated in cells close to the target cell. The results in Fig. 3 use $D = 2000\text{km}$ and $r = 6$. In addition, in the simulation results shown in Fig. 3 we additionally weight cells by the malaria prevalence, as estimated, which makes the simulation more realistic scenario.

#### *1.15.6 Simulation implementation*

We implemented the above model (Equation (S10)) using the WebGPU framework to efficiently carry out computation across many grid cells in parallel using a GPU. In brief, we used a discretised version of our posterior mean estimated HbS frequency map and implemented the simulation in each pixel of this map. We also generated a version of the MAP project 2010 *Pf* prevalence map at the same resolution for use in weighting. A WebGPU compute shader was then written to implement (Equations S1-S3) in parallel across grid cells.

Expression (S10) potentially involves summing over many thousands of nearby grid cells. To make this more computationally efficient, we approximated this by a sum over a fixed number (k) of mosquitoes, each of which chooses a source cell in a uniformly random direction and with distance sampled from the above Beta distribution. In the results presented here we used k=5,000; lower values of k lead to faster but less accurate simulation.

An advantage of the WebGPU approach is that it naturally lends itself to immediate visualisation and interactive exploration of results in a web application framework. We implemented real-time visualisation of genotype frequencies and LD, and a set of controls for fitness and other parameters which behave interactively (in the sense that altering their values takes immediate effect on the simulation.). For custom visualisation work (e.g. Fig. 3) we also implemented an export feature that allows the current set of allele frequencies to be downloaded. A custom R function was written to enable import into R.

#### *1.15.7 Choice of fitness parameters in simulation*

Although the general relative fitness parameter matrix in our simulation has seven free parameters, constraints on reasonable parameters are provided by our geographic analysis and by previously observed data as described in section **1.7** above. In our simulation framework above, the relative risk of infection conferred by HbS against parasites with a particular genotype $g$ is given by the ratio of values in rows of (S8), i.e.:

$$RR_g = \frac{\gamma_{g,S}}{\gamma_{g,A}} \tag{S16}$$

Taking this into account we focussed on simulations with $RR_{--}$ on the order of 1 in 10 to 1 in 100 (reflecting a strong protective effect of HbS against wild-type parasites), and a $RR_{++}$ close to 1 (reflecting little effect of HbS against ++ parasites). In addition, the fact that parasites with *Pfsa+* alleles are observed in large numbers of infections of A individual suggests that the overall fitness cost of + alleles cannot be large; we therefore focussed on simulations where $\gamma_{++,A} > 80\%$. Finally, relative risk estimates suggest that parasites which carry intermediate numbers of + alleles (e.g. -+ or +- in our simulation) may have intermediate levels of fitness, but the exact strength of this is unclear[13] at present. Three specific models we considered are:

*Additive model*: effects in which the major fitness costs add across alleles separately within A and S individuals:

$$\begin{array}{ccc} \gamma_{-+,A} = 1 + s_A & \gamma_{+-,A} = 1 + s_A & \gamma_{++,A} = 1 + 2s_A \\ \gamma_{-+,S} = \gamma_{--,S} + s_S & \gamma_{+-,A} = \gamma_{--,S} + s_S & \gamma_{++,S} = \gamma_{--,S} + 2s_S \end{array} \tag{S17}$$

where in our simulation $s_A < 1$ and $s_S > 1$.

*Multiplicative model*: effects in which the major fitness costs multiply across alleles, separately within A and S individuals:

$$\begin{array}{ccc} \gamma_{-+,A} = s_A & \gamma_{+-,A} = s_A & \gamma_{++,A} = s_A^2 \\ \gamma_{\pm,A} = \gamma_{--,S} \cdot s_S & \gamma_{\pm,S} = \gamma_{--,S} \cdot s_S & \gamma_{++,S} = \gamma_{--,S} \cdot s_S^2 \end{array} \tag{S18}$$

where in our simulation, $0 < s_A < 1$ and $s_S > 1$.

*Dominance model*: fitness cost is incurred by either, or both, of the low-fitness alleles at either loci, separately within A or S individuals:

$$\begin{array}{c} \gamma_{\pm,A} = \gamma_{\pm,A} = \gamma_{++,A} = s_A \\ S\gamma_{-+,S} = \gamma_{+-,S} = \gamma_{--,S} = s_S \end{array} \tag{S19}$$

where in our simulation, $0 < s_S < 1$ and $0 < s_S < 1$.

Specific parameters that appeared to work well in our simulation implementation are:

$$\Gamma = \begin{bmatrix} & A & S \\ -- & 100\% & 1\% \\ -+ & \gamma_{-+,A} & \gamma_{-+,S} \\ +- & \gamma_{+-,A} & \gamma_{+-,S} \\ ++ & 82\% & 82\% \end{bmatrix}, \tag{S20}$$

where the fitnesses of -+ and +- genotypes take one of the forms above.

### 2 Supplementary Figures

#### 2.1 Fig. S1. Predictions of haemoglobin sickle allele frequency in the study area.

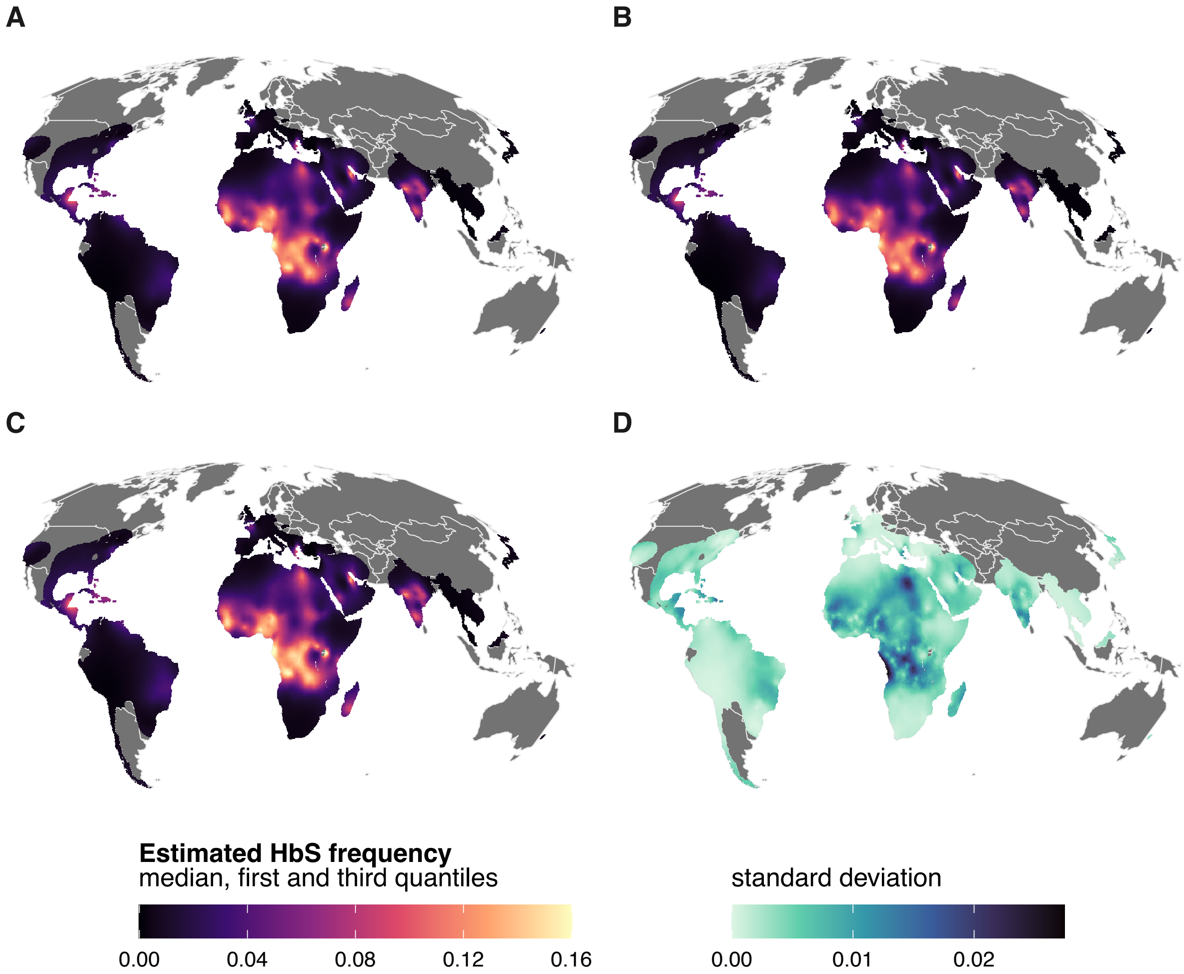

**Fig. S1. | Predictions of haemoglobin sickle allele frequency in the study area.** The median (**A**), first quantile (**B**), third quantile (**C**), and standard deviation (**D**) obtained from 10,000 posterior samples of the haemoglobin sickle (HbS) prevalence model are shown in a 5km x 5km grid in Mollweide projection. Colours in panels A-C show low values in dark purple blue and high values in bright yellow while in panel D, high values are highlighted in dark blue and low values in light green. Predictions are not provided in areas (grey) where HbS is not expected to be present.

2.2 **Fig. S2 Comparison of sickle haemoglobin allele frequency (HbS) to previous work, in hexagonal cells worldwide.**

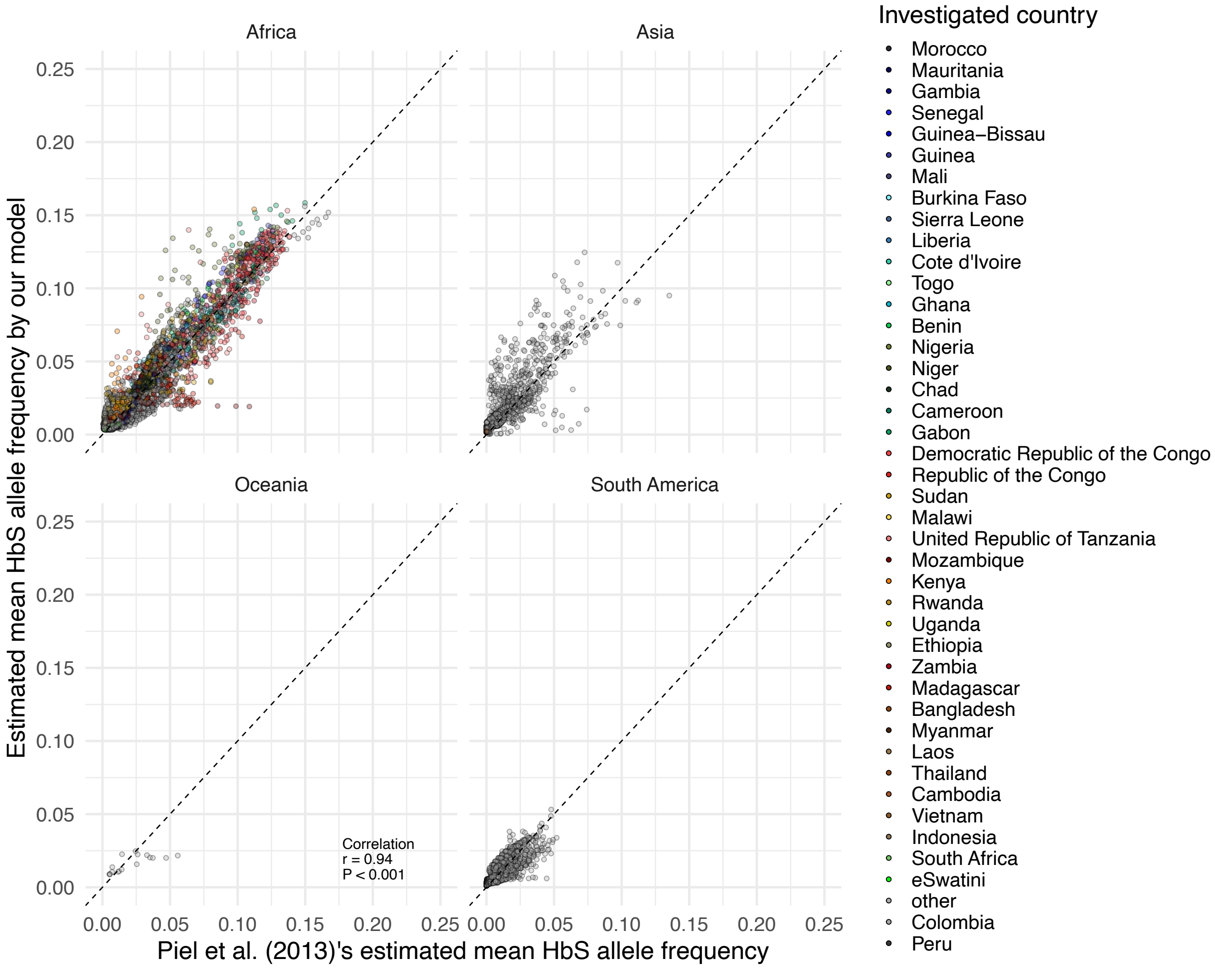

**Fig. S2. | Comparison of sickle haemoglobin allele frequency (HbS) to previous work, in hexagonal cells worldwide.** The mean HbS predicted values based on our model and data (*y-axis*) are compared with HbS predicted values obtained in Piel et al.[3] (*x-axis*) and aggregated at hexagonal level in four regions (panels). Each point represents a hexagon coloured by its corresponding country. The overall correlation given by Pearson's *r* value and *P* representing its associated P-value.

### 2.3 Fig. S3 HbS allele frequency map and association with *Pfsa1*+ allele for different range and standard deviation parameter values of the HbS geospatial regression model.

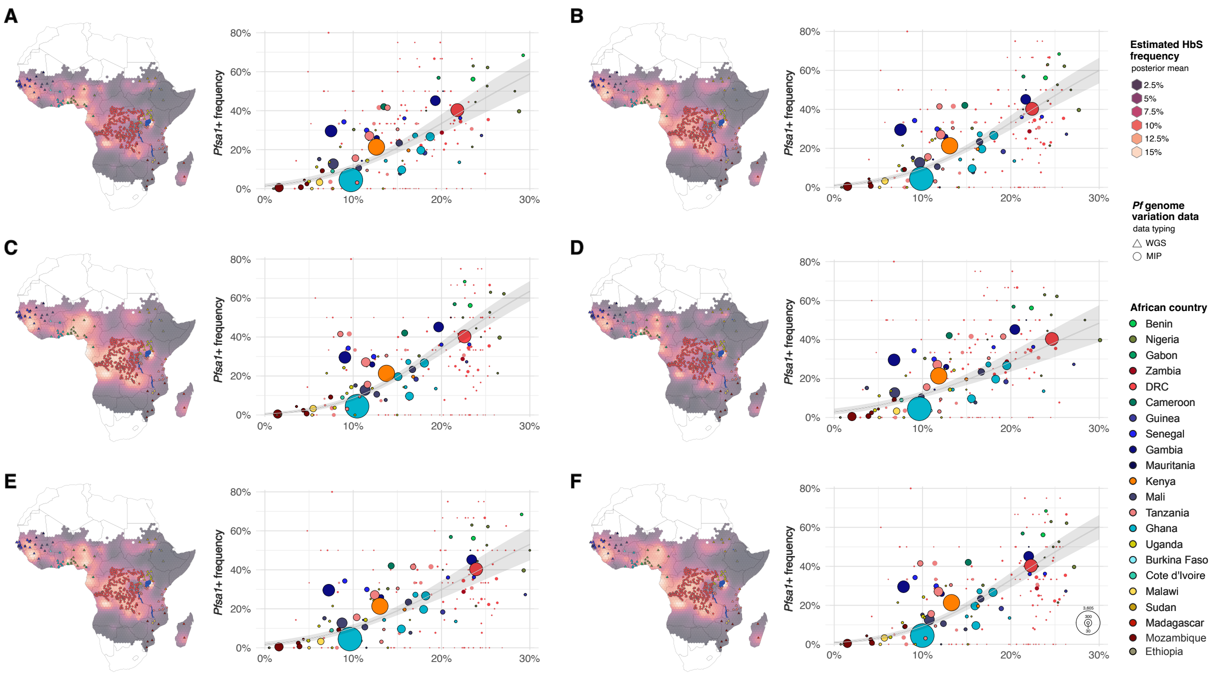

**Fig. S3 HbS allele frequency map and association with *Pfsa1*+ allele for different range and standard deviation parameter values of the HbS geospatial regression model.** This figure shows (left) the estimated HbS mean frequency across hexagons in malaria-endemic African regions (areas with presence of *Pf* parasite observed in 2000 among population of children between 2 and 10 years old) and (right), the association between the combined frequency of HbAS and HbSS genotypes ($f_{HbAS/SS}$, $x$ $axis$) with *Pfsa1*+ ($y$ $axis$). The panels show the results with different values of the range ($r$, in degree) and standard deviation ($\sigma$) parameter **(A)**: ($r = 5$; $\sigma = 0.6$), **(B)**: ($r = 10$; $\sigma = 0.6$), **(C)**: ($r = 25$; $\sigma = 0.6$) which corresponds to the parameters used for the results reported in the manuscript, **(D)**: ($r = 5$; $\sigma = 1$), **(E)**: ($r = 10$; $\sigma = 1$), and **(F)**: ($r = 25$; $\sigma = 1$).

### 2.4 Fig. S4 Association between HbS and *Pfsa1*+ in different regions across Africa.

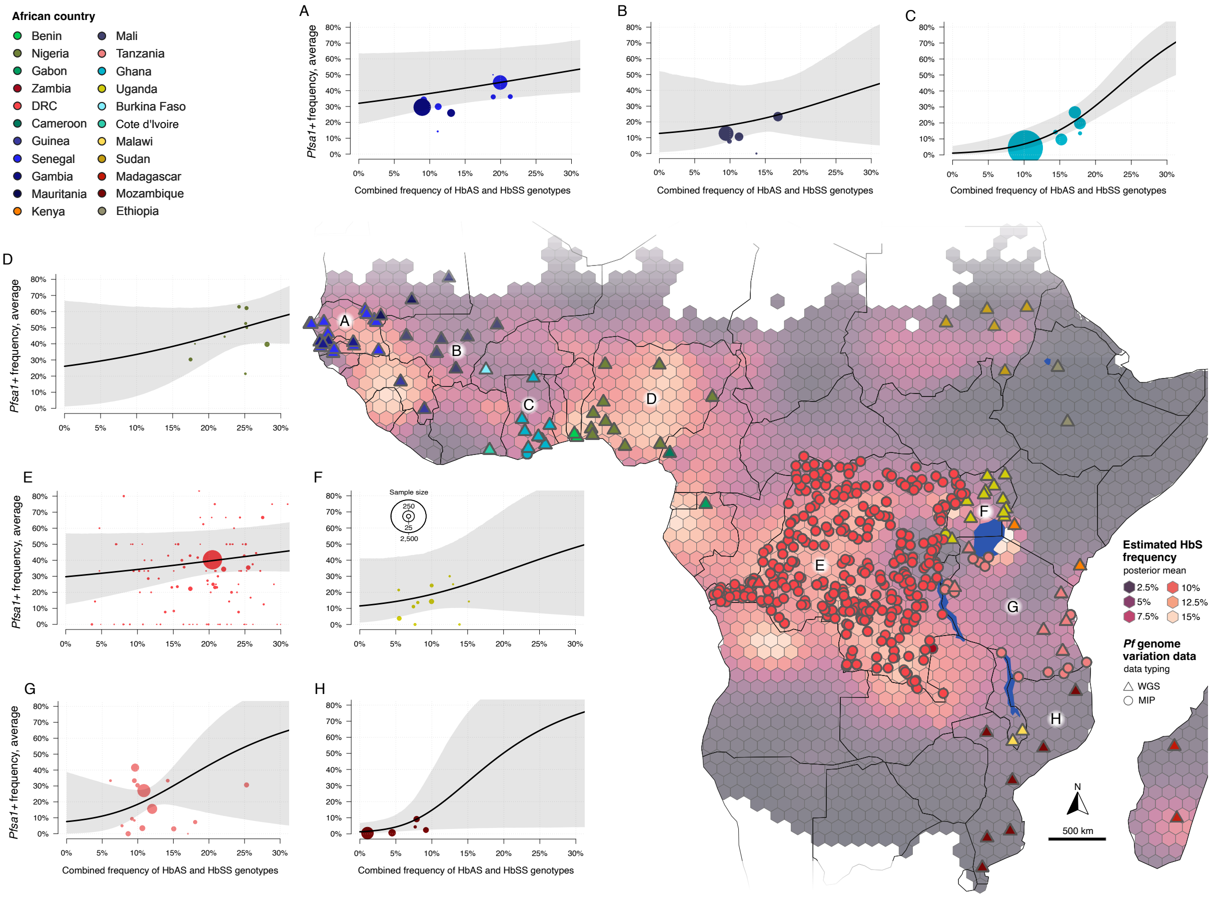

**Fig. S4 | Association between HbS and *Pfsa1*+ in different regions across Africa**. Panels A-H show *Pfsa1*+ allele frequency (*y-axis*) plotted against $f_{HbAS/SS}$ (*x-axis*) at hexagonal level, as in Fig. 1, C and D but estimated using only data from the illustrated regions. The analysis is carried out separately for: (**A**) Senegal and The Gambia, (**B**) Mauritania, (**C**) Ghana, (**D**) Nigeria, (**E**) Democratic Republic of Congo, (**F**) Uganda, (**G**) Tanzania, and (**H**) Mozambique. The African map highlights the estimated HbS mean frequency across hexagons in malaria-endemic regions (areas with presence of *Pf* parasite observed in 2000 among population of children between 2 and 10 years old).

### 2.5 Fig. S5 Detail of long-range linkage disequilibrium between *Pfsa*+ alleles.

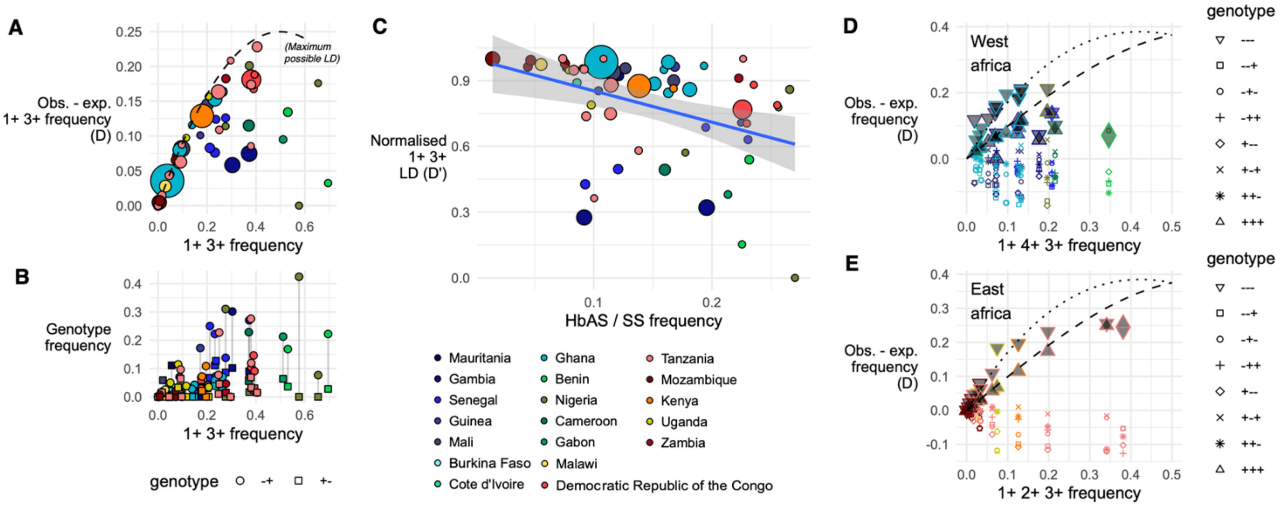

**Fig S5 | Detail of long-range linkage disequilibrium between *Pfsa*+ alleles.** (**A**) Linkage disequilibrium (LD; computed as Lewontin's D, *y axis*) between *Pfsa1*+ and *Pfsa3*+ alleles, plotted against the *Pfsa1/3* ++ allele frequency (*x axis*) in each hexagon across Africa. Only hexagons containing at least 50 Pf data points are shown. The dashed line represents the highest possible value of D given the *x axis* value. (**B**) the frequency of combined -/+ and +/- genotypes at the *Pfsa1* and *Pfsa3* loci in each hexagon in Africa. (**C**) Normalised LD (D') plotted against HbAS/SS genotype frequency across Africa. (**D**) and (**E**) 3-way LD metric (similar to panel (A)) for each possible genotype at the *Pfsa1*, *Pfsa4*, and *Pfsa3* loci in west Africa (panel D) and the *Pfsa1*, *Pfsa2*, and *Pfsa3* loci in east Africa (Panel E). The combined genotypes are shown by shapes, with the +++ and −−− genotypes shaded as in the legend. The dashed line shows the maximum possible LD for the +++ genotype given its frequency (similar to panel A), while the dotted line shows the LD for the −−− genotype that would be seen if it were the only other genotype present.

### 2.6 Fig. S6 Theoretical constraints on parasite fitness

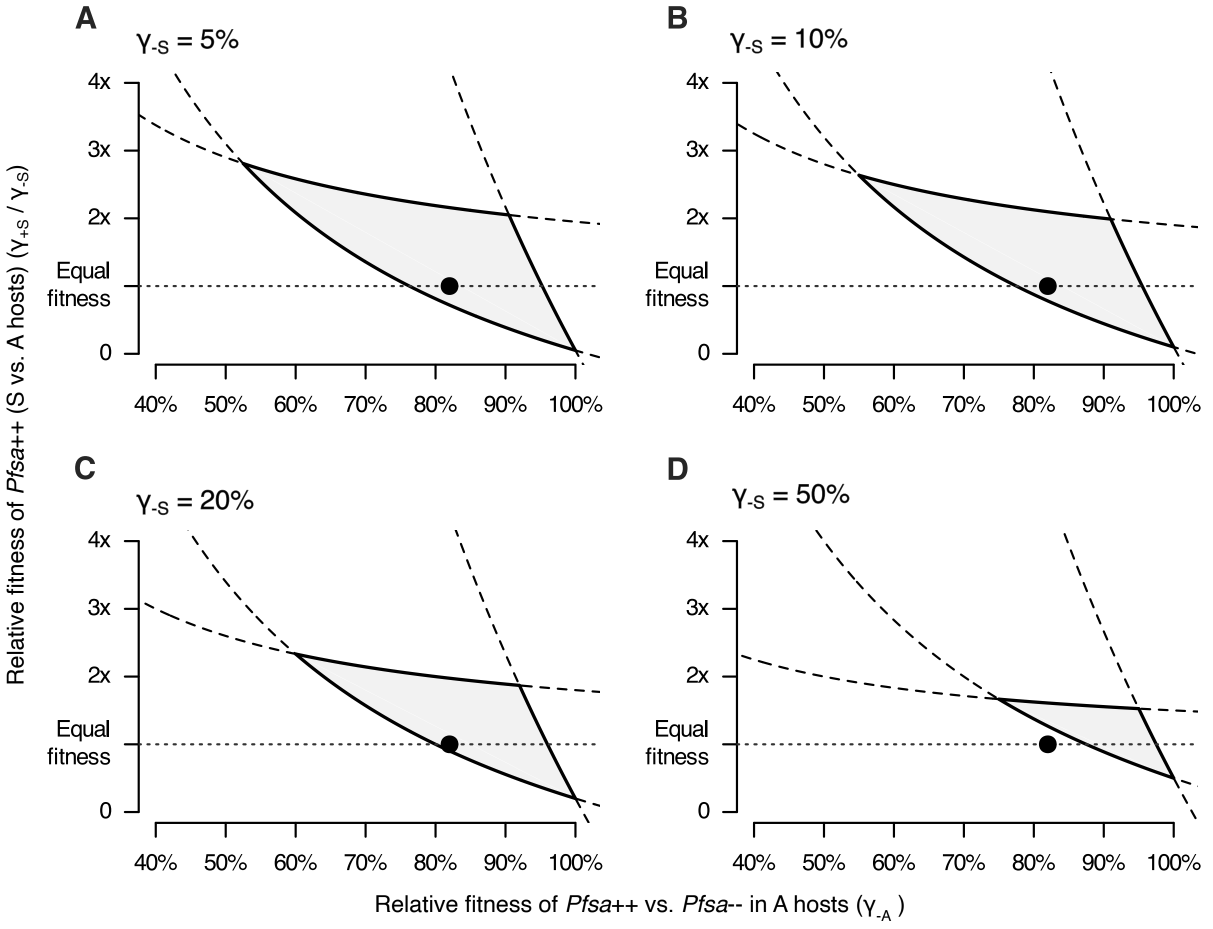

**Fig S6 | Theoretical constraints on parasite fitness.** Plot shows plausible values of parasite fitnesses (grey area) under assumptions described in Methods, and assuming that *Pfsa*++ parasites are not positively selected at $f_{HbAS/SS} < 5\%$ and not negatively selected at $f_{HbAS/SS} > 25\%$ (lines $l_1$ and $l_2$), and that HbS does not confer increased parasite fitness even when the *Pfsa*+ is at high frequency ($f_{++} = 50\%$, line $l_3$). The results depend on the degree to which HbS impedes *Pfsa*- parasites ($\gamma_{-S}$) as shown in the panel title; Fig. 3A shows the constraint when $\gamma_{-S} = 1\%$. The dot indicates values similar to those used in the simulation (Fig. 3B).

### 2.7 Fig. S7 *Pfsa1*+ allele age estimate using GEVA-like method

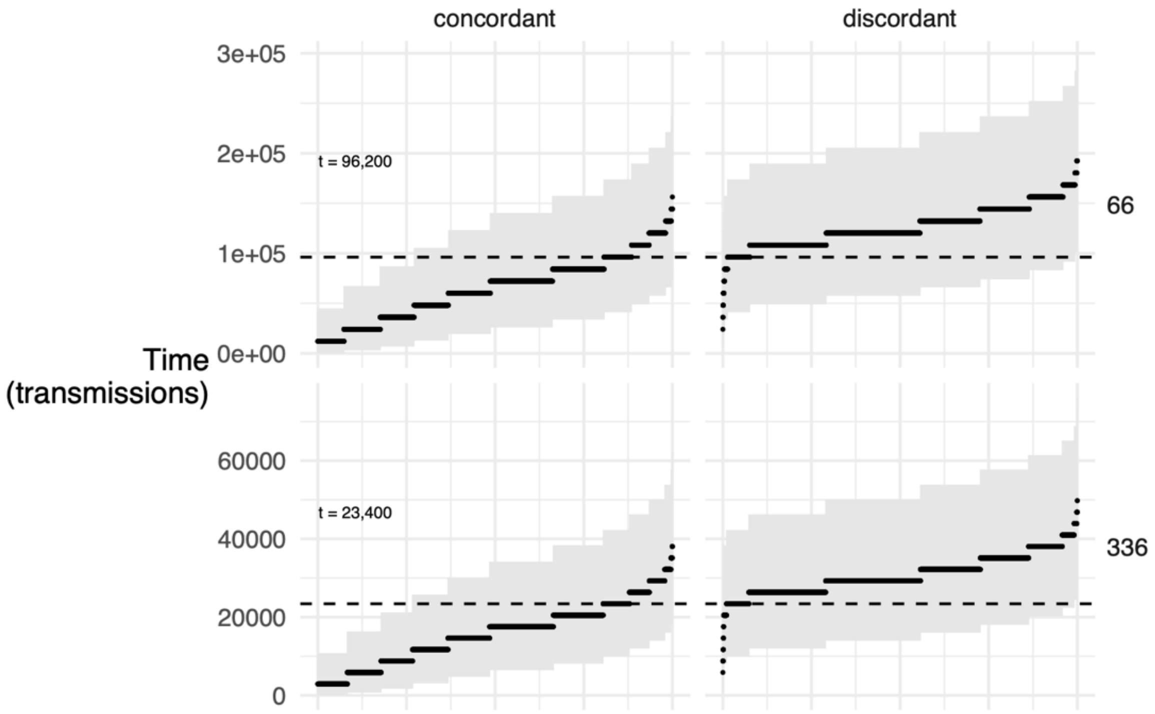

**Fig S7 | *Pfsa1*+ allele age estimate using GEVA-like method.** Plot shows an estimate of the age of the *Pfsa1*+ allele (dashed horizontal lines, in transmission cycles) computed using a GEVA-like method[28], based on comparison of pairwise nucleotide differences between parasites among the N=4,788 from the Pf7 popgen dataset. Estimates are calculated using variation over a 5kb window centred at the *Pfsa1*+ lead variants (chr2:631,190 T>A). Left panels: posterior mean (black points) and 95% posterior interval (grey areas) of the time to most recent common ancestor for a random subset of 10,000 pairs of parasites which both carry the *Pfsa1*+ mutation ("concordant" pairs, x axis). These estimates provide a lower bound for the age of the allele. Right panels: the same information for a random subset of 10,000 pairs of parasites in which one carries *Pfsa1*+ and the other *Pfsa-* ("discordant" pairs). These estimates provide an upper bound for the age of the allele. The dashed line and text show the estimated age, chosen as the value which minimises the number of pairs above (left panels) or below (right panels) the line. The two panel rows correspond to an assumption of 66 and 336 mitoses per generation respectively, and the per-base, per-mitosis mutation rate is assumed to be $3.83 \times 10^{-8}$; these values are chosen as in the calculation in Otto et al[23]. An effective population size of 50,000 was assumed. This method is adjusted from the GEVA method[28].

### 2.8 Fig. S8 Haplotype patterns and estimated genealogy at the *Pfsa3* locus.

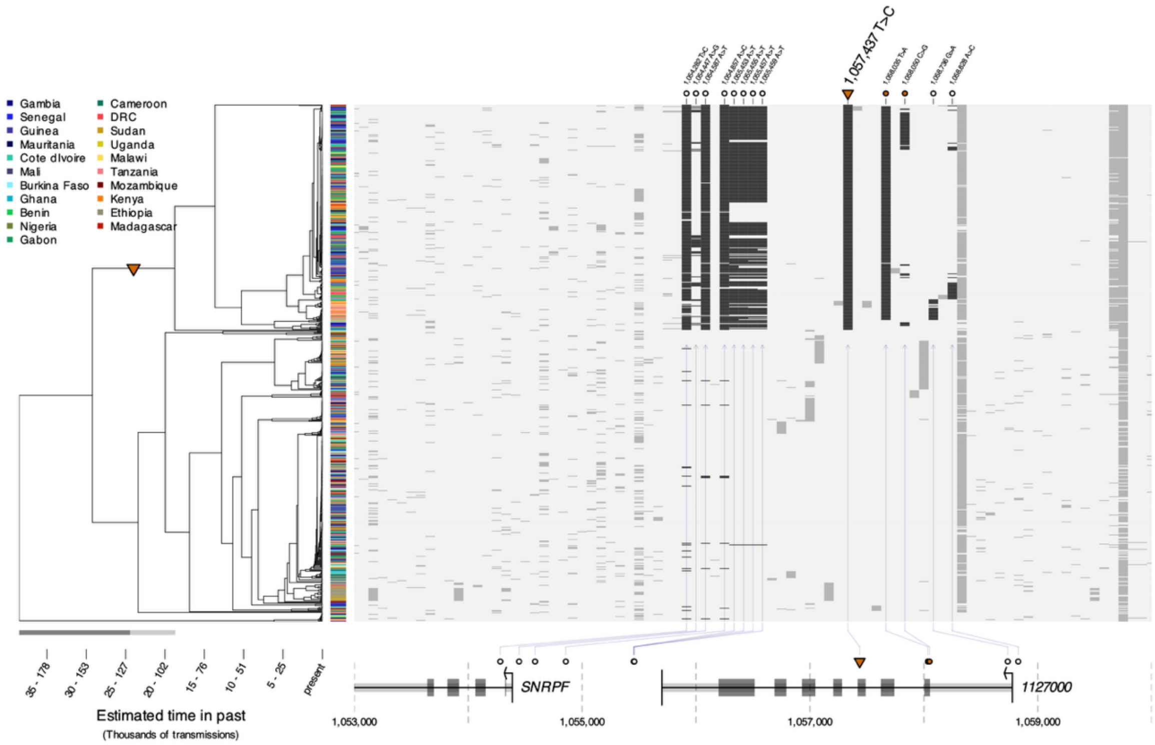

**Fig S8 | Haplotype patterns and estimated genealogy at the *Pfsa3* locus.** This analysis suggests that the chr11:1,057,437 T>C allele (which was second most strongly associated with HbS in the analysis of severe cases[13]) is most ancestral, and we focus the figure on this allele. Figure details are as in Fig. 4 legend. The highlighted variants are those with less than 5% frequency on *Pfsa3*- background, at least 5-fold higher frequency on the *Pfsa3*+ background, and at least 2% frequency overall.

### 2.9 Fig. S9 Longitudinal estimates of *Pfsa1*+ allele frequencies

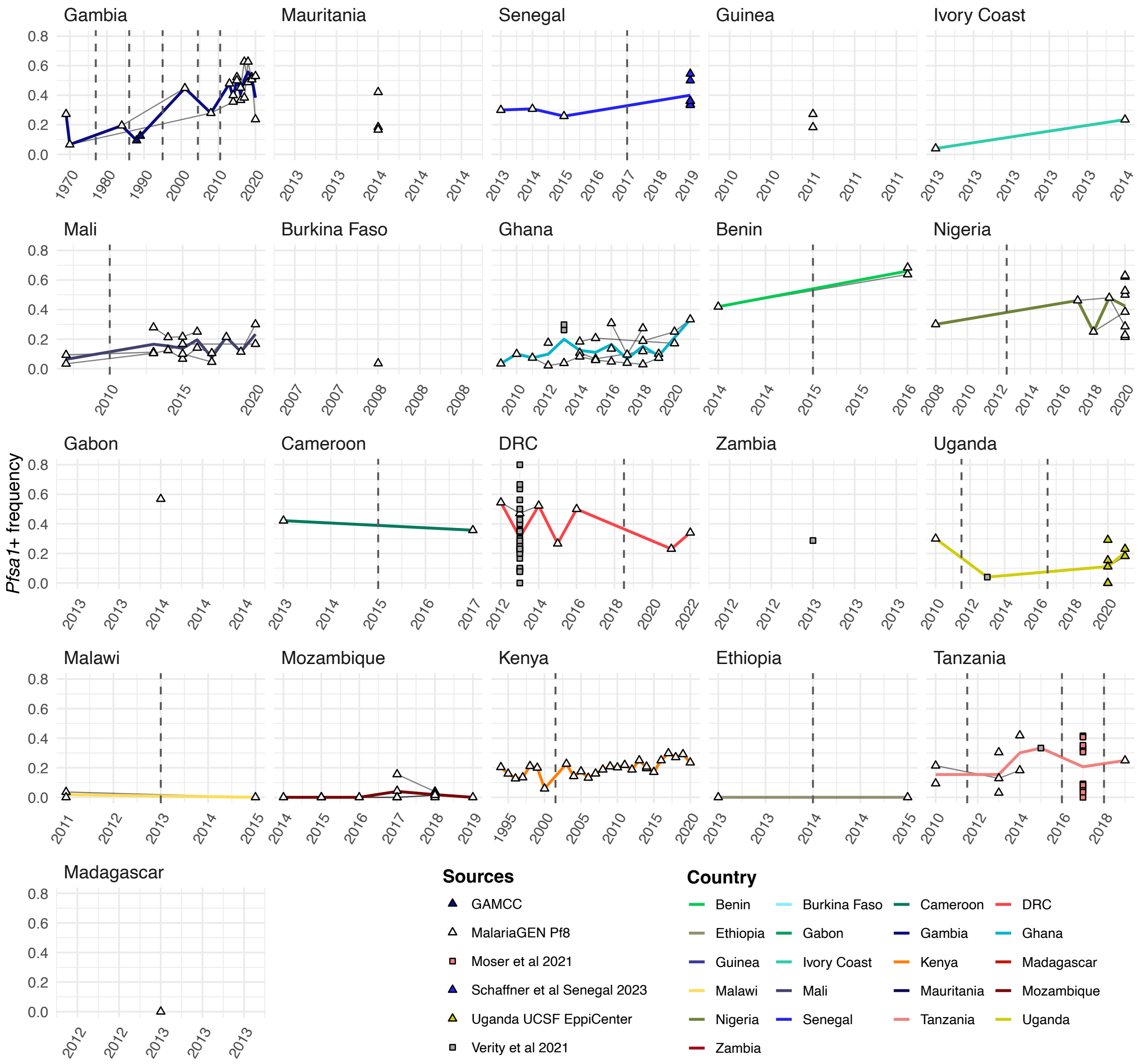

**Fig. S9 | Longitudinal estimates of *Pfsa1*+ allele frequencies.** Points show data estimates of *Pfsa1*+ allele frequency in hexagons (as in Fig. 1) in African countries with at least one site including more than 10 samples. Panels are sorted longitudinally. Data sources are indicated by coloured shapes with triangles that denote WGS, and squares for MIP data typing. Estimates in the same geographical location (hexagon) in each country are joined by grey lines. Yearly averages by hexagon are linked with a thick line coloured by country. A vertical dashed line indicates temporal gaps where more than one year of data collection is missing. Slopes intersected by dashed lines should be therefore interpreted with caution, as they may not reflect an underlying trend.

### 2.10 Fig. S10 Longitudinal estimates of *Pfsa3*+ allele frequencies

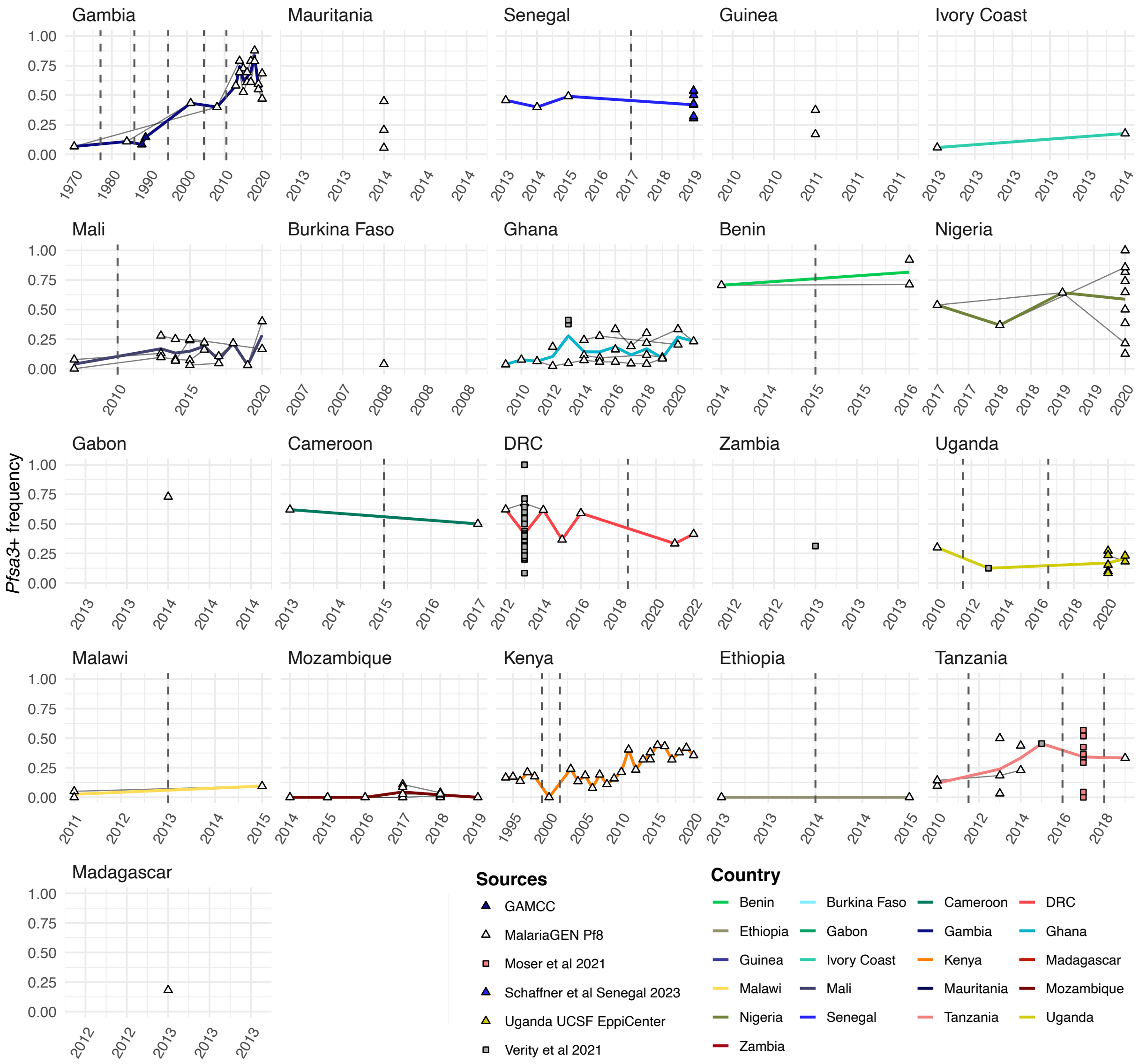

**Fig. S10 | Longitudinal estimates of *Pfsa3*+ allele frequencies.** (**A**) Points show data estimates of *Pfsa3*+ allele frequency in hexagons (as in Fig. 1) in African countries with at least one site including more than 10 samples. Figure details are as in Fig. S9 legend.
